# Benchmarking tools for DNA repeat identification in diverse genomes

**DOI:** 10.1101/2021.09.10.459798

**Authors:** Gourab Das, Indira Ghosh

**Affiliations:** School of Computational & Integrative Sciences (SC&IS), Jawaharlal Nehru University (JNU), New Mehrauli Road, New Delhi 110067, Delhi, India; Army Hospital Research & Referral, Near Military Hospital Road, Subroto Park, Dhaula Kuan, New Delhi 110010, Delhi, India

**Keywords:** DNA Repeats, Tool comparison method, Genome analysis, Benchmarking, Web-server

## Abstract

Continuous progression in genomics shows that repeats are important elements of genomes that perform many regulatory and other functions. Eventually, to date, many computational tools have been developed and frequently used for the identification and analysis of genomic repeats. A single tool cannot detect all different types of repeats in diverse species rather pipeline of tools is more effective. But, the choice of such rigorous and robust tools is highly challenging. A method has been implemented to select a set of optimal tools for finding all available classes of perfect and imperfect tandem repeats including microsatellites, minisatellites, and interspersed CRISPRs in genomes. A total of 11 tools have been shortlisted using rule-based selection and then ranked by analyzing rigorousness in searching in diverse species and execution time. Tool comparison shows consistency in perfect microsatellite detection performance but significantly differ for long and imperfect repeats. A web-server has been built which provides a generic platform for various classes of repeat identification from the diverse genome using multiple tools and comparison.

## 1. Introduction

From small viral genomes to large plant genomes, the occurrence of repeats has been observed everywhere. In small prokaryotic genomes like bacteria, repeats are less abundant and mostly found inside genes. But in large eukaryotic genomes e.g. human, plants, they are ubiquitous in the non-coding region covering a major fraction of the genome [1–3]. For example, there exist very little or statistically no differences among great ape genome sizes (Human: 2.88 Gbp, Chimpanzee: 2.99 Gbp, Gorilla: 3.08 Gbp and Orangutan: 3.04 Gbp) but the number of genes (Human: ∼33k and other primates: ∼55k with 95% human orthologs) and STRs (short tandem repeats) are significantly different [4]. According to their presence, they perform a wide variety of functions including genetic and epigenetic regulation, genome packaging and stability maintenance, binding sites of transcription factors, structuring protein topology, and many more [5]. Moreover, they serve as mutational hotspots in the genome [6–8]. In phenotypic context, repeats assist in bacterial adaptation and pathogenesis, the cause of many genetic disorders in humans, help plants to survive under stressful conditions [9–11]. Till now, numerous classes and subclasses of repeats are detected and classified in many ways like, tandem and interspersed according to the arrangement of repeating motifs; based on motif lengths, tandem repeats (TRs) can be further divided into microsatellites or simple sequence repeats (SSRs) or short tandem repeats (STRs) of motif length 1-6 bp and minisatellites which include variable number tandem repeats (VNTRs) are of motif length 7-100 bp. Depending on the nature of repeating motifs, again tandem repeats can be perfect or imperfect, exact or approximate, etc. Interspersed repeat constitutes of many families and sub-families of transposons or transposable elements (TEs). Using the underlying mechanism of transposition, TEs can be divided into two main classes: type I or retrotransposons and type II or DNA transposons. Understanding repeat’s diversity, ubiquity in genomes and functional importance, many computational tools have been developed using a variety of algorithms to detect and analyze these repeats. Successive reviews have discussed the basic protocol and underlying algorithms for repeat identification [12–15] which have been summarized in Figure 1.

**Figure 1:**
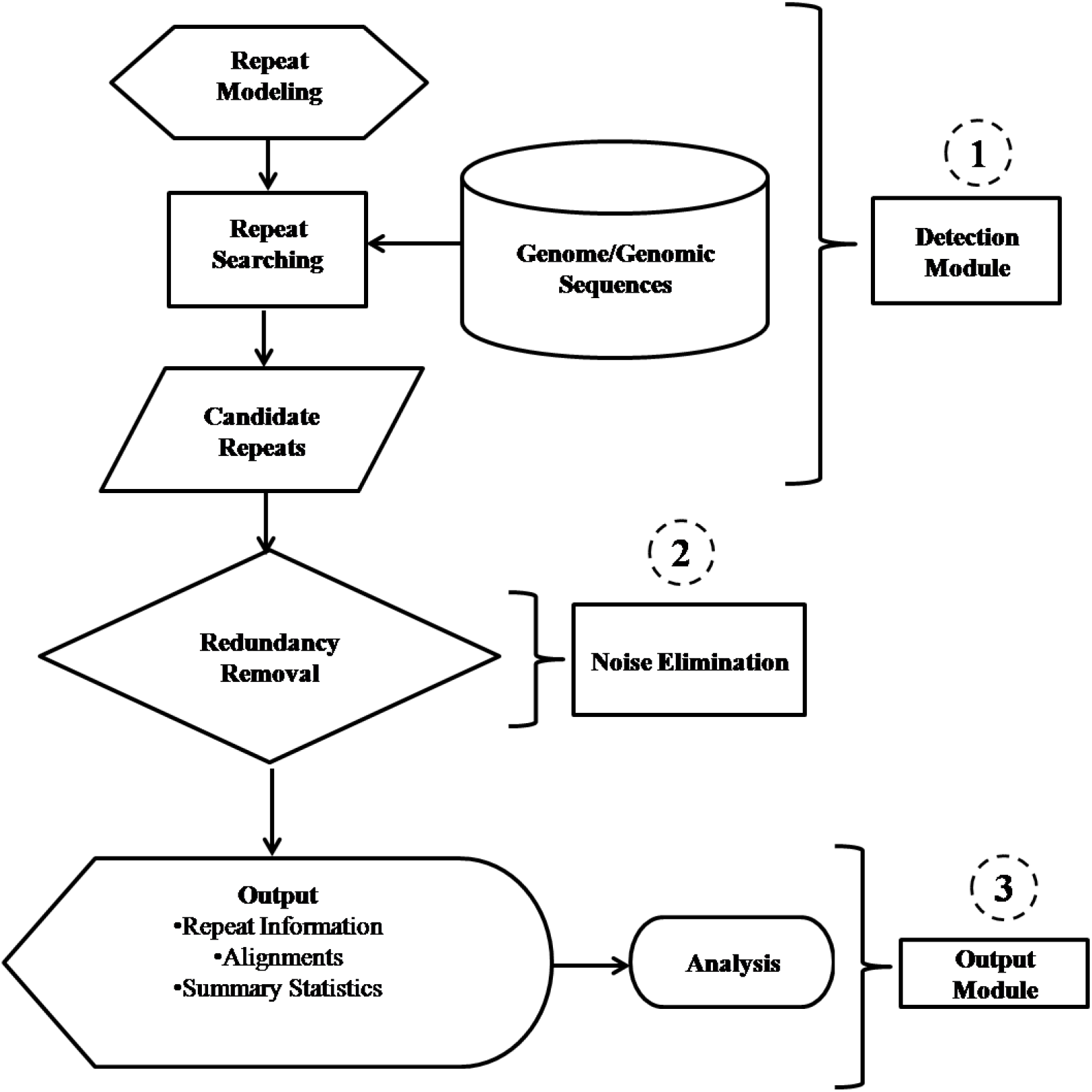
Basic modules of repeat detect protocol. (1) represents the detection part; (2) stands for quality control or noise removal module; (3) is the output generating unit followed by downstream analysis.

Repeat finding tools made up of three distinct modules to perform their work: the First module is for repeat detection. To search for repeats in the genomic/genomes, different tools have followed many modeling approaches to build repetitive patterns [16,17]. These include a combinatorial method, regular expression, *k-mer* sliding window-based approaches. Next, finding these patterns in the genomes has been accomplished either by exhaustively or by heuristically to report a list candidate repeats. Exhaustive search is found to be comparatively slower than heuristic or probabilistic search but former technique ensures rigorousness in repeat identification. To select candidate repeats, several statistical criteria have been applied to detect the plausible repetitive sequences [12,13,18,19]. In the second module, after building and searching the candidate repeats, redundancy in the candidate list has been checked and a unique one has been selected on basis of several user-defined cut-offs to produce the final list of repeats. In the third and last module, repeat statistics have been calculated; output style has been formatted and reported. Following this basic protocol, several tools have been designed and upgraded continuously to identify the aforementioned classes of repeats. The earlier review has suggested building a pipeline of tools for efficient detection of all classes of repeats present in genome [16]. But due to the availability of plenty of tools listed in the earlier reviews, the choice of the best performer remains a difficult task.

The current study addresses this problem and has developed a method to rank the tools so that the pipeline can produce a comprehensive list of repeats from the input genome. To test the robustness of the method, it has been applied to the genomes from diverse branches of the tree of life with distinct properties. Following the protocol (Figure 1) and using the criteria like rigorousness, robustness, execution time, etc. tools have ranked and results have been presented. Additionally, a web-server RDTcs has been built to provide a common platform for various repeat detection tool comparison from genomes of user’s interest. Furthermore, an exclusive list of repeat detection tools has been presented which can make the existing list [20] more comprehensive (Table 1).

**Table 1:**
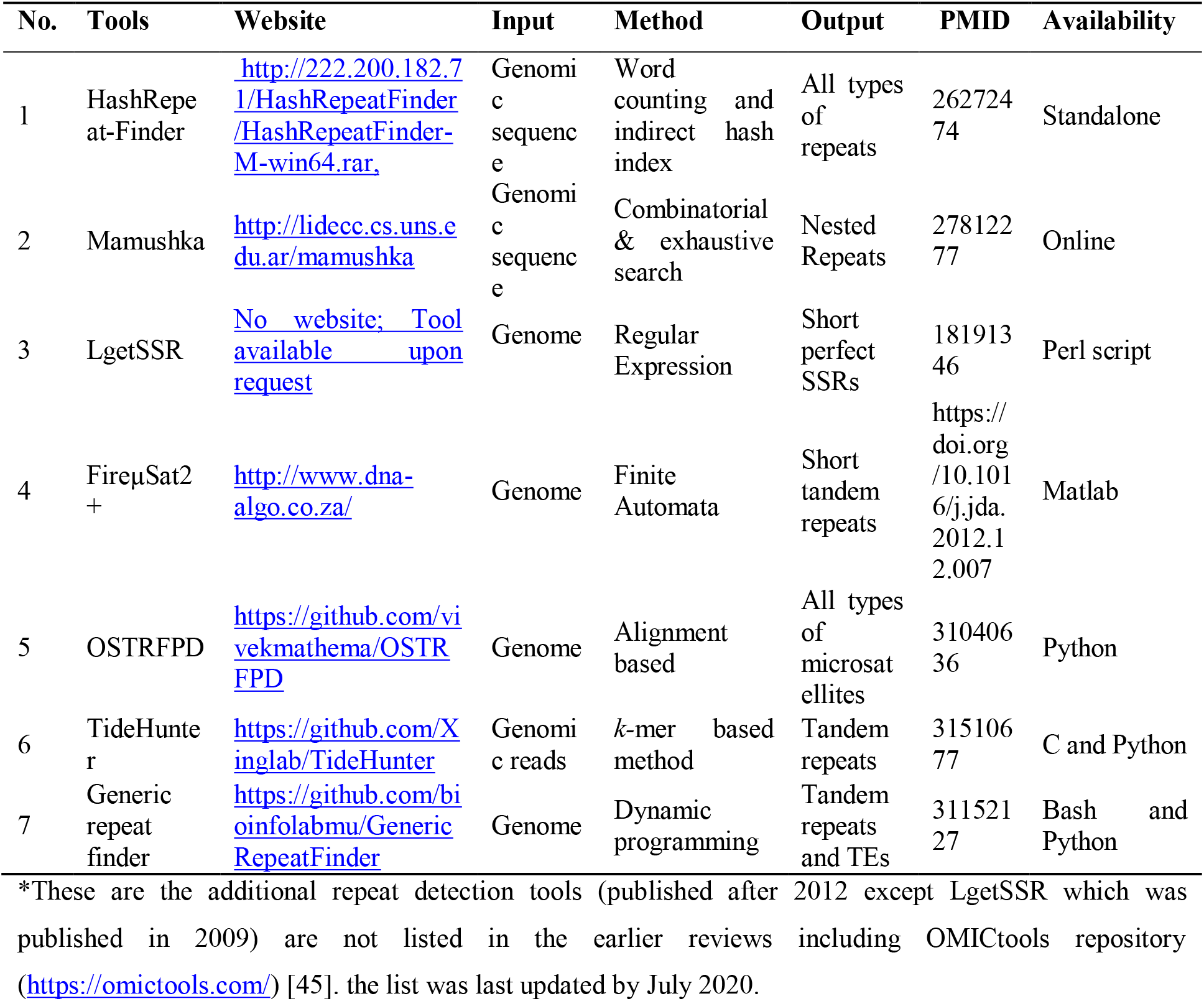
An exclusive list of different types of repeat identification tools*

## 2. Materials and Methods

### 2.1 Genome Dataset

Genomes are chosen very selectively for the extraction of repeats and comparison of tools. Different biological systems comprising of prokaryotes like viroid, virus, bacteria, and eukaryotes like protozoa, plant have been included in the list and their fasta sequences have been downloaded from RefSeq release 76 [21]. All of these systems are quite different from each other in context to different properties. For example, bacterial genomes are comparatively small, intron-less, and contain a single chromosome only whereas plants are larger, possess multiple chromosomes with introns. Inside the bacterial community, there exist many differences among the genomes. Some are AT or GC rich whereas some are neutral like *E. coli*. Some are gram-positive in contrast to some gram-negative bacterial strains which are also included in the list. Again both pathogenic and non-pathogenic systems have been opted to enrich the diversity of the selected dataset. So, tools that are compatible and efficient with these different systems have been listed for rigorousness testing. Different strains and species with varying genomic properties have been selected to test the robustness. Genome details of the aforementioned systems are given in Table 2. Genome assemblies of prokaryotes are complete whereas gaps are present in the eukaryotic genomes.

**Table 2:** List of selected genomes used in the present study

| No. | Organism | Kingdom/ Group | RefSeq Assembly / Accession | Size | %GC | Remarks |
| --- | --- | --- | --- | --- | --- | --- |
| 1 | *Escherichia coli* str. K-12 MG1655 | Bacteria | GCA_000005845; NC_000913 | 4.64 Mbp | 50.8 | Non-pathogen; Gram -ve |
| 2 | *Mycobacterium tuberculosis* H37Rv | Bacteria | GCA_000195955; NC_000962 | 4.41 Mbp | 65.60 | Pathogen; Gram +ve |
| 3 | *Bacillus subtilis* str. 168 | Bacteria | GCA_00000904; NC_000964 | 4.42 Mbp | 43.50 | Non-pathogen; Gram +ve |
| 4 | *Shigella flexneri* 2a str. 2457T | Bacteria | GCA_000007405; NC_004741 | 4.60 Mbp | 50.9 | Pathogen; Gram -ve |
| 5 | *Anaeromyxobacter* *dehalogenans* 2CP-1 | Bacteria | GCA_00002214; NC_011891 | 5.03 Mbp | 74.70 | Non-pathogen; Gram -ve |
| 6 | halophilic archaeon DL31 | Archaea | GCA_000224475; NC_015954 | 3.64 Mbp | 62.38 | Unclassified |
| 7 | *Plasmodium falciparum* 3D7 | Protozoa | GCA_000002765 | 22.9 Mbp | 19.04 | Malaria parasite |
| 8 | Amsacta moorei entomopoxvirus | Virus | NC_002520 | 232.39 Kbp | 19.00 | Bio-controlling agent of pests |
| 9 | Coconut cadang-cadang | Viroid | NC_001462 | 0.246 Kbp | 59.30 | Plant Pathogen |
| 10 | Avocado sunblotch viroid | Viroid | NC_001410 | 0.247 Kbp | 38.10 | Plant Pathogen |
| 11 | *Arabidopsis thaliana* | Plant | GCA_000001735 | 119.67 Mbp | 36.05 | Eudicot |
| 12 | *Brachypodium distachyon* | Plant | GCA_000005505 | 272.13 Mbp | 46.44 | Monocot |

### 2.2 Selection of tools

Tools have been shortlisted for further comparisons using the following criteria: first, tools must handle small to large genomes; second, if tools are capable of detecting more than one class of repeats, then it must have separate option for each class of repeats unlike Tandem Repeat Finder (TRF) [22]; third, ability to generate user-friendly output with at least start-end coordinates and repeating motif information included; fourth, availability of standalone command-line versions for running in local computer in batches; fifth, do not account contiguous ‘N’ letters as repeats; sixth, compatible with single sequence and multi-fasta format and seventh, can identify novel repeats. However, for the tools where multi-fasta and user-friendly output options are not satisfied like Tandem Repeat Occurrence Locator (TROLL) [23], output formatting is managed by the inclusion of custom shell scripts for further post-processing. To search repeats, TROLL uses a microsatellite motif (length 1-6 bp) dictionary having all possible combinations of ‘A’, ‘T’, ‘G’, ‘C’ including mono- to hexameric repeats completing the perfect microsatellite repeat space. Hence, searching using this tool can identify all perfect microsatellites in the genome. But, post-processing scripts will increase the total execution time of repeat extraction. On the other hand, TRF is dedicated to finding long imperfect tandem repeats efficiently and used as the fundamental tool for extracting imperfect tandem repeats in the pipeline programs and several databases like TRDB [24]. Finally, a set of tools has been selected using the rules mentioned above (Table 3).

**Table 3:**
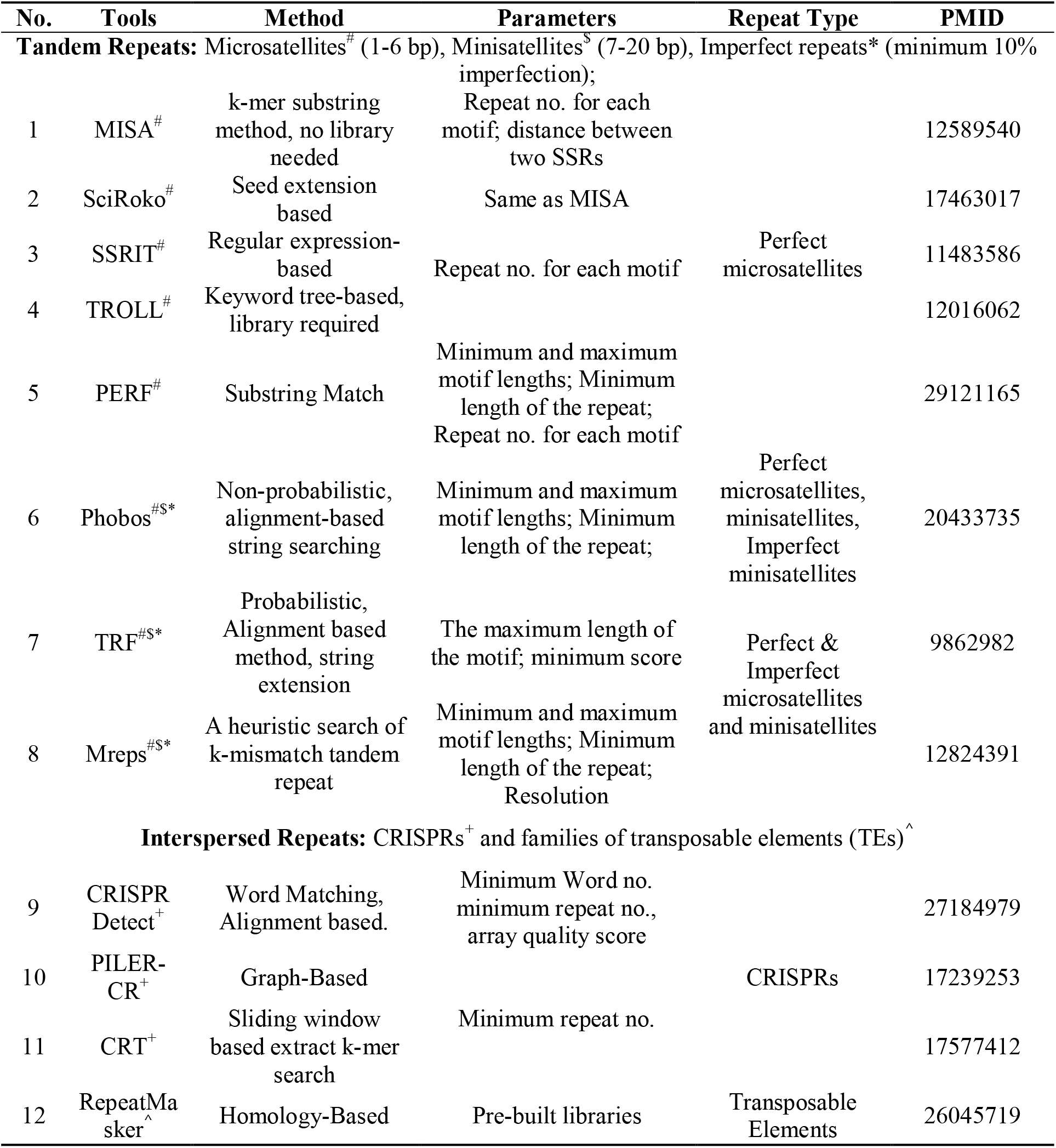
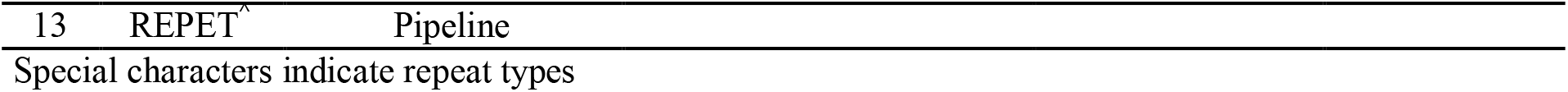
Tool details with input parameters and output repeat types

### 2.3 Parameters setting and repeat extraction

One important and tricky step is to tune the parameters of the tools so that it can find all possible repetitive loci in a particular genome. As already stated, many tools based on several algorithms of computational science, are available which can identify various types of repetitive patterns in the genome. Naturally, default parameter settings used by the tools are very different from each other. For example, TRF has minimum repeat length cut-off 50 bp whereas MISA [25,26] uses 10 as repeating copies for minimum cut-off. It is very confusing to set comparable/equivalent parameter values for all the tools for the identification of a specific class of repeat. For perfect repeat, it is observed that minimum repeat no. (no. of repetitions) and repeat length is the common parameters in most of the tools. These two parameters are interchangeably used as one can be derived from the other if repeating motif length is known. In the case of imperfect repeats, other such common parameters are minimum repeat length, minimum score the percentage of imperfection. But, the main challenge remains in the determination of cut-offs for an imperfect repeat. That is why earlier reviews [12] have suggested relying on the default parameters because that must have been benchmarked and tested for the identification of plausible repeat candidates. For perfect microsatellites, the length of the repeats is possibly determined by several mutational events like strand slippage, homologous recombination, etc. It is also suggested that length of microsatellite loci can be 12 for mono-to tri- and 16 for tetra-, 20 for penta- and 24 for hexameric motifs [27] but these values may vary from species to species. This set of values has been used in this work for microsatellite extraction. As TROLL and Phobos [28], both do not have any minimum repeat no. input option, hence custom shell-scripts have been written so that it can take repeat no. as input too. All other tools like MISA, SSRIT [29], SciRoko [30], PERF-SSR [31] can take minimum repeat no. as an input parameter. Following Castelo et al. MISA, SSRIT, SciRoko, PERF-SSR minimum repeat no. has been set as follows: 30 for mono-, 15 for di-, 10 tri- and 8 for tetra-, 6 for penta- and 5 for hexameric motifs. Beier et al. have used repeat no. set 10-6-5-5-5-5 for mono-di-tri-tetra-penta-hexameric motifs [23,32]. Another parameter like the distance between two SSRs in MISA has been set to zero. Except for the parameter sets mentioned in Table 3, all other parameters for these programs have been fixed by default.

In the case of perfect minisatellites, only two tools are preferred i.e. TRF and Phobos. Minimum repeat length is one of the suitable parameters that can be used for comparison. TRF does not have the minimum repeat length input parameter instead minimum score has been calculated by multiplying match weight of Phobos has the default input minimum repeat length parameter. For minisatellites motif length ranging from 7-20 bp, minimum repeat length 40, 60, and 80 bp have been used. In the case of imperfect repeats, both minimum repeat length and minimum repeat no. have been used along with an additional 10% imperfection for detecting imperfect repeats. While using repeat no. the parameter, set of values suggested by Merkel et al., has been utilized for imperfect microsatellite detection. For imperfect minisatellites of motif lengths ranging from 7-20 bp, minimum repeat no. is chosen as 2. The minimum repeat length parameter has been set as 40 and 60 bp.

There are several parameters used by CRISPR detection tools like repeating motif lengths, no. of repetitions, length of the spacers, length of the flanking regions on both sides, etc. Default values for most of these parameters have been fixed by using the available knowledge on CRISPRs which has come from the experiments. Here only no. of repetitions have been varied and set as 2, 3, and 4 for comparing CRISPR detection tools. For each class of repeats, different parameters and species have been used to incorporate randomness and robustness for comparing tools in an unbiased manner.

### 2.4 Repeat density-based tool’s evaluation

Two different measures are widely used to characterize repeats i.e. repeat density in number (*D*_*N*_) and repeat density in length (*D*_*L*_). For a particular genome, *D*_*N*_ is defined as a total no. of repeats per Mbp of the genome and *D*_*L*_ is the total length in base pair covered by per bp of the genome. These two parameters signify two different events i.e. trade-off between duplication-deletion events and tolerance level of a genome respectively. These repeat densities have been calculated as done by Achaz et al.[33]

## 3. Results

### 3.1 Testing correctness of tool’s configurations

A total of 11 tools has been selected based on the rules described in section 2.2 for comparing the abundance of repeats in the genomes (Supplementary Table T1). Tools have been configured in a 1600 MHz processors cluster with 8GB Mb RAM having CentOS 6.0 operating system. To check whether tools have been properly installed or not, results from the literature where the particular tool’s usage has been observed are taken to reproduce the result using the same parameter settings and genome/genomic data. Supplementary Table T1 summarizes the reproducibility of published work results and inspects the correctness of the tool’s configuration in the local server.

### 3.2 Comparison of tools with different parameters in each repeat class

Using the parameters for each class of tool as mentioned in Table 3, three different sets of values have been selected to extract repeats from the genomes listed in Table 2. These diverse sets of values of the parameters have been taken from the published articles and used for repeat detection to check the consistency in tool performance. Tools have been compared with respect to repeat density parameters to test rigorousness in repeat detection, execution time, and robustness with different datasets and parameter sets.

#### 3.2.1 Check for rigorousness

Comparisons of repeat density parameters for 6 selected perfect microsatellites detection tools have been shown in Figure 2 (*D*_*N*_) and Supplementary Figure F1 (*D*_*L*_) which depicts that all of the perfect microsatellite detection tools are performing almost equally for prokaryotic systems and comparable in case of eukaryotic genomes except *Plasmodium falciparum*. Three tools with names TROLL, Phobos, and PERF_SSR are equally efficient in the detection of maximum no. of repeats per Mbp in each of the genomes as compared with SciRoko and MISA which are little poorer in performance.

**Figure 2:**
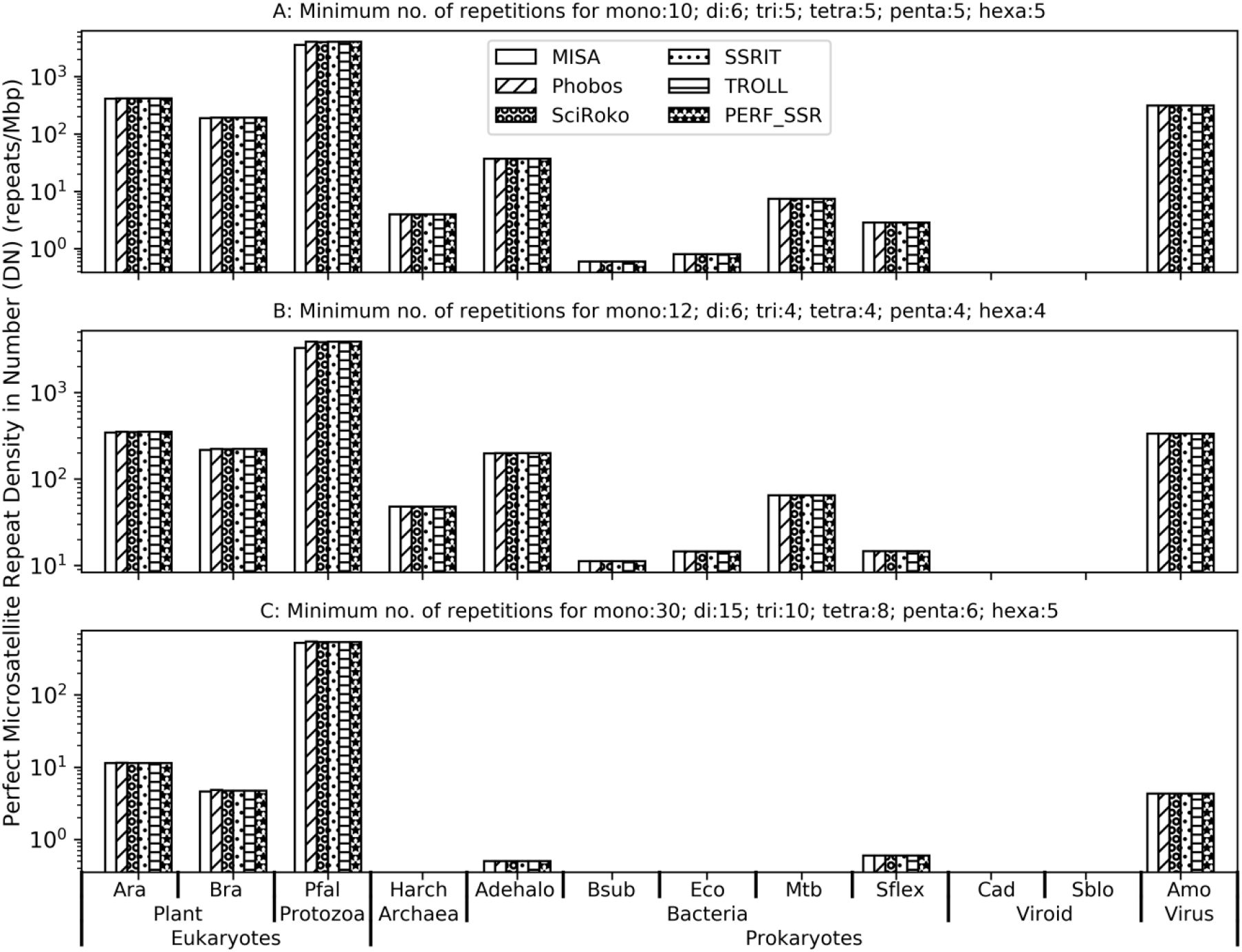
Comparison of repeat density in number parameter (*D*_*N*_) of perfect microsatellite detection tools using a diverse set of genomes and 3 distinct set of values for repeat no. parameter. Y-axes are on a log scale.

Rigorousness testing result for perfect minisatellite detection tools has been shown in Figure 3 (*D*_*N*_) and supplementary Figure F2 (*D*_*L*_). Phobos has identified the maximum no. of perfect minisatellites per Mbp than TRF from all the genomes. By default, TRF reports both perfect and imperfect minisatellites. The final list of perfect minisatellites has been prepared by filtering out the imperfect ones from TRF output. Using three different values of the Minimum repeat length parameter, repeats have been recorded by Phobos and TRF. Consistent results have been observed which marks Phobos as the more rigorous tool between the two for all parameters and genomes except Amsactamoorei entomopoxvirus where both tools have performed equally well. In the case of imperfect repeats, as shown in Figure 4, Mreps [34] has identified maximum no. of repeats per Mbp and maximum percentage in length (Supplementary Figure F3) of each genome analyzed in this study in comparison to the other tools. A similar result has been observed in earlier publications also [12].

**Figure 3:**
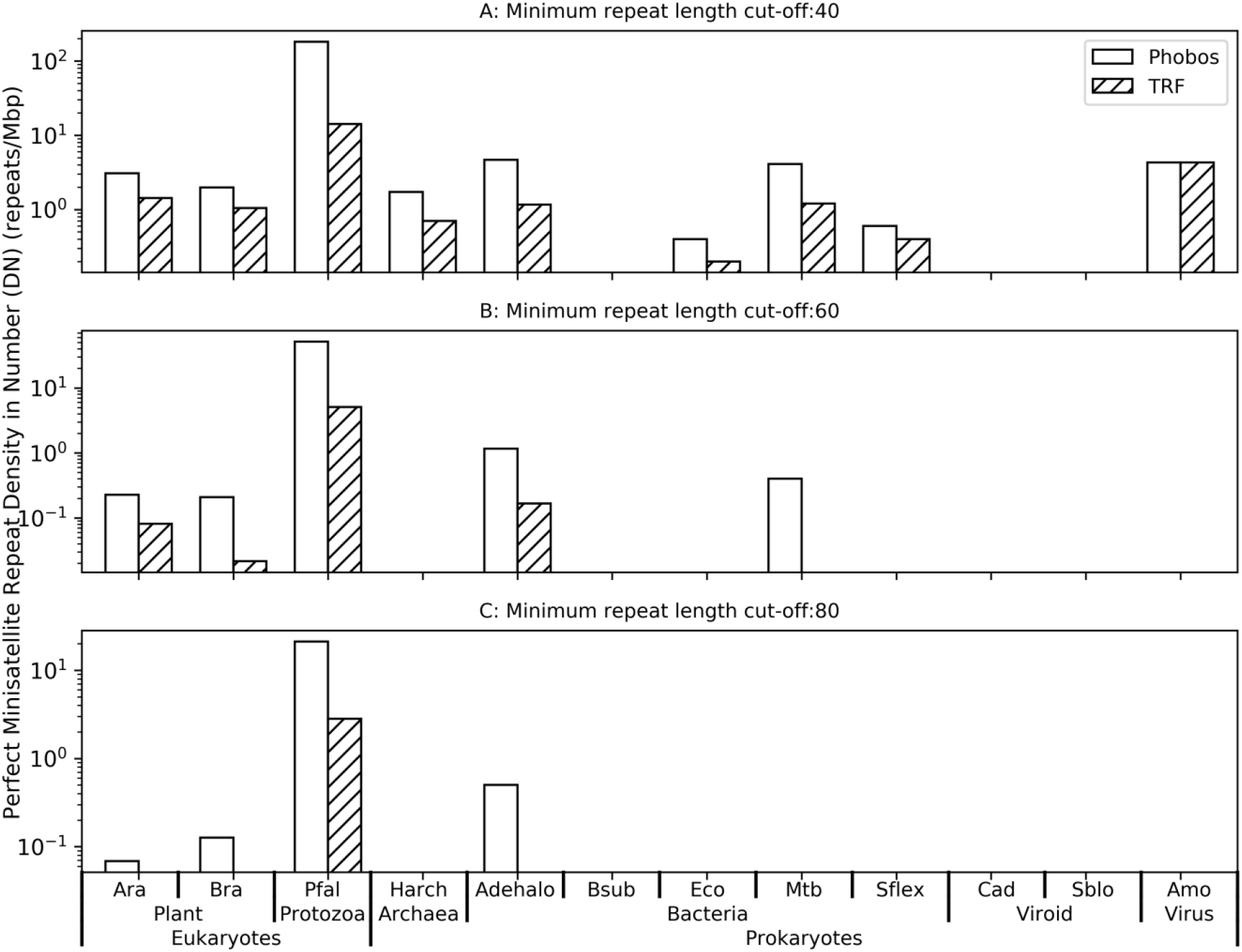
Comparison of repeat density in number parameter (*D*_*N*_) of perfect minisatellite detection tools using a diverse set of genomes and 3 distinct sets of values for repeat no. parameter. Y-axes are on a log scale.

**Figure 4:**
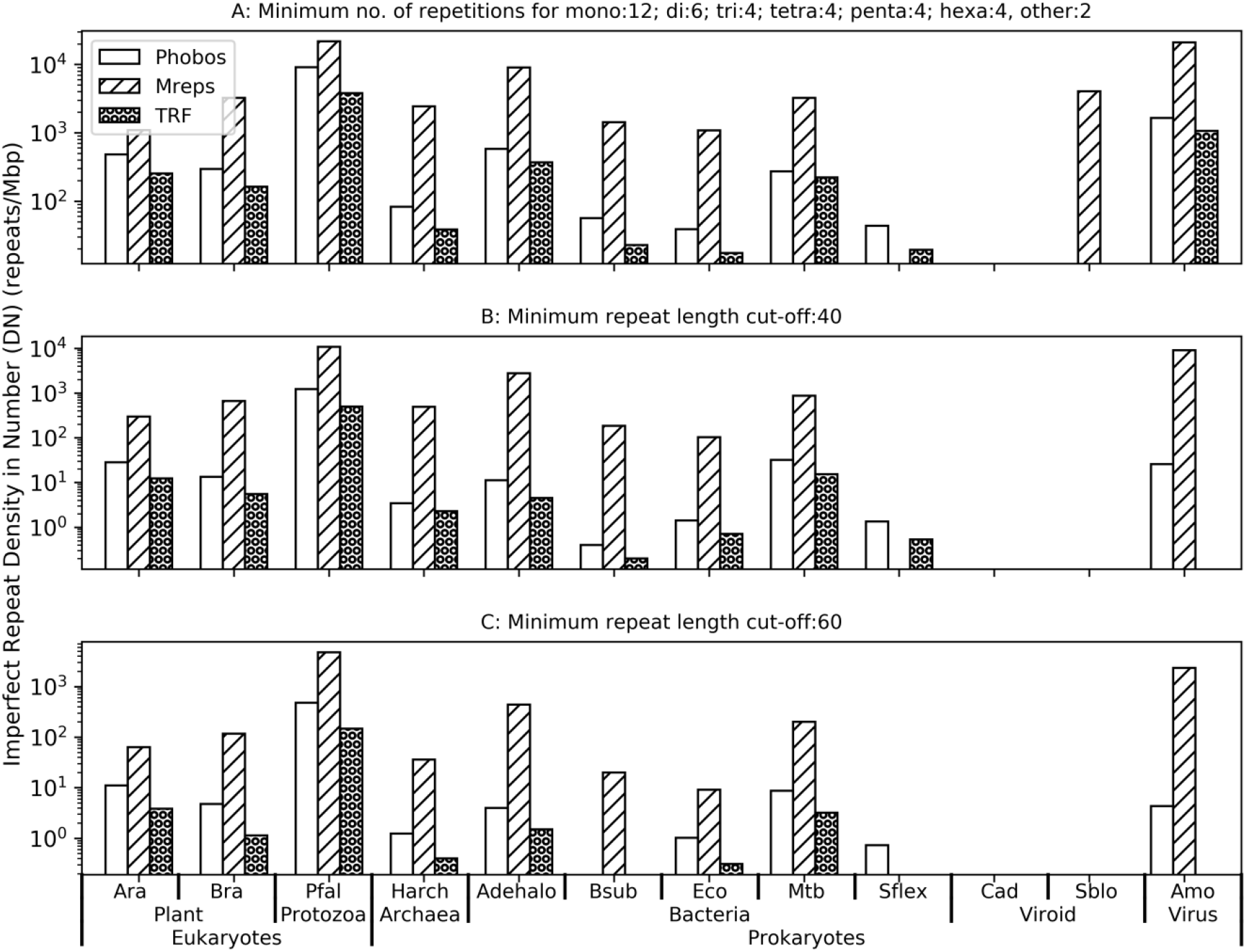
Comparison of repeat density in number parameter (*D*_*N*_) of imperfect tandem repeat detection tools using a diverse set of genomes and 3 distinct sets of values for repeat no. parameter. Y-axes are on a log scale.

A comparison of the tool’s performance in CRISPRs detection has been performed among three standalone tools. CRISPR has a specific sequence and structural organization in the genome and also presents very few in numbers. The definition of CRISPRs has been nearly standardized regarding the length of the repeat, length of the spaces, lengths of the flanking region, etc and tools have been developed following the definition. Using three different values of repeat no., tools have been compared. Among the three tools, CRT [35] has found a maximum no. of CRISPRs per Mbp (Figure 5) and also in percentage length (Supplementary Figure F4) of the genome followed by CRISPR-Detect [36] and Piler-CR [37]. Both of these tools have performed comparably except Anaeromyxobacter dehalogenans where CRISPR-Detect performs better than Piler-CR. But, CRISPR-Detect fails in detecting palindromic sequences in the eukaryotic genomes. CRISPRs are one kind of interspersed repeats found in prokaryotic genomes mainly. But eukaryotic genomes are full of several interspersed repeat classes like Retrotransposons, DNA transposons, etc. Several tools have been developed to identify these interspersed repeat classes and successfully reviewed recently [18,38]. However, the database ‘Repbase’ [39]repbase has been designed for storing manually curetted interspersed repeat classes. Two tools namely RepeatMasker [40] and REPET [41] have been employed for the extraction of interspersed repeats from various eukaryotic genomes. A comparison has been done between pre-built libraries. Results show that RepeatMasker has identified a total of 45491 interspersed repeats including several families and sub-families of DNA and retrotransposons covering more than 98% of the unique repeats detected by the two tools. Only 692 (1.5%) repeats have been uniquely found in REPET (RepBase20.05) library. The majority (∼74%) of the repeats have been recognized by both tools (Supplementary Figure F5).

**Figure 5:**
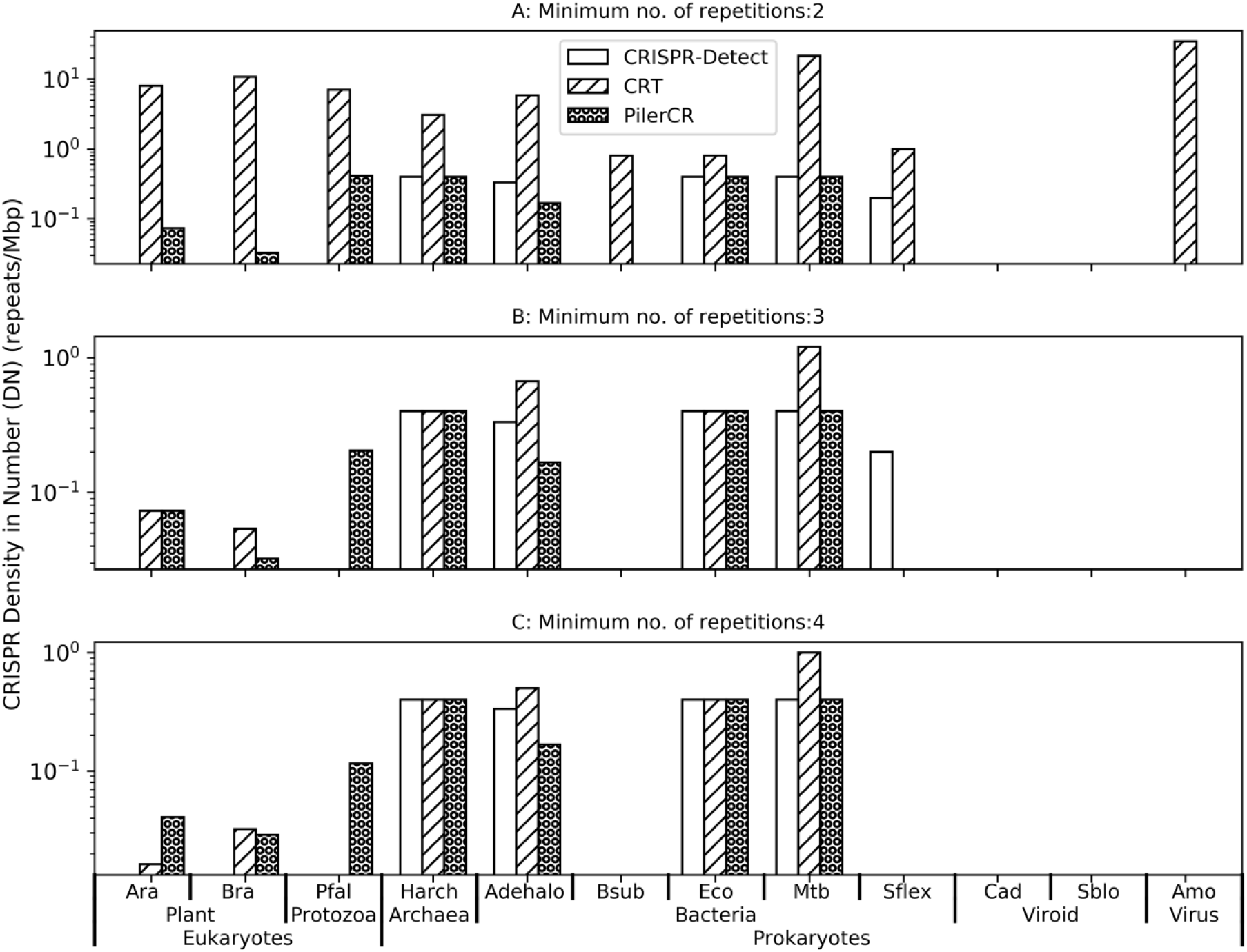
Comparison of repeat density in number parameter (*D*_*N*_) of CRISPR detection tools using genomes and 3 distinct sets of values for repeat no. parameter. Y-axes are on a log scale.

#### 3.2.2 Comparing execution time

Despite comparable performance in rigorous detection of perfect microsatellite repeats in diverse genomes, tools significantly differ in execution time in large genomes. In the case of small genomes (up to 5 Mbp genome length), all tool’s execution times are almost similar (Supplementary Figure F6 and F7) but for larger genomes, SciRoko is the fastest and MISA seemed to be slowest among the tools. Searching long microsatellites (of length 30 bp long or more) Phobos performs better than Sciroko. The order of other tool’s time of execution remains the same.

In terms of execution speed of perfect minisatellite tools, Phobos is far superior to TRF except for the small prokaryotic genomes where both tool’s execution times are comparable. For larger eukaryotic genomes, Phobos is approximately 4 times faster than TRF (Supplementary Figure F8 and F9). The reason behind the slow behavior of TRF is the filtering module using statistical measurements. On the other hand, Phobos reports all possible high scored alignments including overlapping and non-overlapping repeats without any filtering treatment which reduces the execution time.

While comparing execution time of the imperfect repeat programs, similar observation has been found like micro- and minisatellites i.e. for larger genomes like plants, high variations in execution times have been found whereas for small-sized genomes results are comparable (Supplementary Figure F10 and F11). Repeat detection times for Mreps and TRF are comparable except minimum repeat no. parameter values where Mreps found to be the fastest tool followed by TRF and Phobos has been marked as the slowest tool. This slow speed problem of Phobos has been notified by the developer in the manual which stated that in imperfect search of motif lengths greater than 10 bp Phobos will run slower than expected.

All of the three tools for CRISPR identification namely CRISPR-Detect, CRT, and Piler-CR have detected CRISPRs incomparable time for prokaryotic genomes. In the case of eukaryotes, CRT and Piler-CR have shown the same efficiency in searching time as prokaryotes (Supplementary Figure F12-F13). Long execution time for eukaryotes indicates the same fact about CRISPR-Detect that due to refinement using several filtering modules exhausts the time.

#### 3.2.3 Testing robustness of the tools

The robustness of the tools can be checked in many contexts. One way is to tune the program parameters with different values and checking whether the tool’s performance has been changed or not. In the context of input sequences, the tool’s robustness can be also checked. Using input sequences from diverse species with highly variable genomic properties like genome length, %GC content, gram staining, and phenomic properties like pathogenicity, cotyledon structures of plant seed, etc. robustness can be also tested. Using sequences with diverse properties and checking whether tools have been performing similarly for all the sequences or not, the robustness of the tools can be easily examined.

In the case of perfect microsatellite repeats, all of the tools have accomplished repeat detection with similar efficiency but varied in execution speed. This holds for all the parameter values selected in the work for perfect microsatellite identification. While checking the perfect minisatellites tool’s robustness, though both TRF and Phobos can deal with input sequences of different types of organisms with diverse genomic and phenomic properties and behave similarly for a different set of values of the parameters. For each parameter set, Phobos works better than TRF both in terms of rigorousness in repeat extraction and execution time. Regarding imperfect repeat tools, all of the tools i.e. Mreps, Phobos, and TRF are handy with input sequences from any kind of organism including prokaryotes and eukaryotes and respond similarly to all of the parameter values. CRISPR is found only in bacteria and archaea and all of the CRISPR identification tools can handle any type of prokaryotic genome sequences of any length and acted in the same way for different values of parameters. Additionally, several palindromic sequences have been also identified in eukaryotic genomes pointing out their robustness in working with any kind of sequence. Interspersed repeats in the RepBase have been already listed for different eukaryotic genomes using REPET and RepeatMasker indicating the robustness of the tools having capabilities of dealing with any genome sequences. Except for TROLL, all other tools can manage multi-fasta with more than one sequence in a single file. For TROLL, custom shell script has been written to make TROLL working with multi-fasta too. Another important aspect regarding the tool’s compatibility with next-generation sequences should be addressed. Many of the existing genome sequence assemblies (for both pro- and eukaryotes) has not been finished yet. Moreover, many newly sequenced genomes have been deposited in the databases regularly. So, it is expected that most of these sequences will be formatted as multi-fasta (Contigs, Scaffolds) and contain many ‘N’s in the assemblies. The aforementioned tools have been designed in such a way that it will account for continuous chains of ‘N’s and filter them out. It is noted that many new tools (listed in Table 1) are also coming which are dedicated to next-generation sequences only.

### 3.3 Searching genomic locations and annotations for the repeats

Searching genomic locations and annotations of the repeats is mandatory to enlighten the functional implications of the repeats. The functional roles of the repeats are highly dependent on their locations in the genome. Repeats which are present in the intergenic regions might have some effect on the expression of the associated genes (both in upstream and downstream). Those which are predominately occur inside genes (genic repeats) have different kinds of function affecting the gene-products i.e. transcripts and proteins. It is expected that genic repeats have a direct influence on the sequences, structures, and functions of the mRNAs (transcripts) and proteins. Many examples already have been cited about the roles of genic and intergenic repeats [42]. Most of the tools can report the genomic coordinates for the start and end positions of the repeats in the genome except TROLL. But, measuring the abundance of the repeats in the genome is a little tricky due to the presence of overlapping repeats. Hence, the use of applications like intersectBed and mergeBed from BEDTools suite [43] is essential for correct estimation of the repeat content. Regarding annotations of the repeats, annotations of the genes which contain the repeat are useful to assign the annotation for the same repeat. In the case of intergenic repeats, both upstream and downstream gene annotations can be used as the annotations for the repeats. As the tools are reporting overlapping repeats, hence multiple annotations for the same repeat are also possible.

### 3.4 Development of webserver

To make this benchmarking strategy easily usable and accessible, RDTcs web-server (http://rdtcs.indirag.org/) has been developed. Salient features and user manual can be found in the aforesaid website where instructions for how to use the server have been pictorially depicted.

## 4. Discussion

Tool’s performances have been compared for a total of 5 different (perfect micro- and minisatellites, imperfect, CRISPRs, and TEs) types of repeats. Different parameter sets have been used to check the consistency in the tool’s efficiency. The correctness of tool installation has been checked by reproducing the published results. For analyzing perfect microsatellites detected using 6 tools, to overcome the difficulty in the visualization of 6 sets, the UpSet technique of R package UpsetR [44] has been used which helps to visualize the intersections in a matrix format. Along Y-axis, all available intersections have been plotted. In the X-axis, sets associated with each intersection have been marked along with a bar plot presenting no. of elements in each set (Supplementary Figure F14). Phobos, SciRoko, and PERF have identified more no. of unique repeats than the rest of the tools which have comparable set sizes. The reason is the variation in marking the start-end coordinates of the repeats or edge effects which are disappeared when only repeating motifs are considered for generating the sets (Supplementary Figure F15).

Analysis of perfect minisatellites shows that the majority of the unique repeats (∼90%) belong to Phobos whereas only 5% of the repeats uniquely identified by TRF (Supplementary Figure F16). To check how much edge effects influence repeat prediction, unique motifs identified by the tools have been compared and found that almost all of the motifs are identified by Phobos (∼99%). Only 2 motifs have been uniquely detected by TRF (Supplementary Figure F17). These results emphasize that edge effects do influence repeat extraction. To recapitulate the edge effect, it is described as the differences in repeat boundaries (start-end coordinates) reported by the tools developed using different algorithms.

Due to its algorithmic nature, Mreps turn out as one of the widely used tools for imperfect repeat detection. But the main problem with Mreps is its over-prediction of many degenerate repeats where no. of mismatches is too high to mimic the biological models. Another limitation in Mreps is that it does not report the repeating motif. Analysis of detected imperfect repeats has been conducted between Phobos and TRF only because Mreps doesn’t report repeating motif by default. As a result, finding imperfect motifs by manual inspection is laborious and too much time-consuming work. Standardizing repeats seems to be the most difficult task. Different tools have been built upon different algorithms and expected to detect different types of imperfect motifs. Comparing repeats using exact string matching may not be sufficient for comparison of repeats. Supplementary Figure F18 has shown that only ∼2% of the total unique repeats are common between two tools indicating diversity in the reported repeat lists. Like perfect minisatellites, Phobos has detected maximum unique repeats (76%) in comparison to TRF (22%) and superior in performance hence selected. While considering only motifs, intersection size increased to 8% (Supplementary Figure F19). These results suggest that a consensus between imperfect repeat from different tools is difficult to achieve. Future work is required to optimally map one dataset into another.

Among the tools for CRISPR identification, CRT doesn’t report the consensus representative repeating motif for CRISPR hence not included in the analysis. Both CRISPR-Detect and Piler-CR have detected all unique repeats with no common repeats between them (Supplementary Figure F20). In the comparison of motifs, the majority of them (86%) have been uniquely identified by Piler-CR (Supplementary Figure F21). Due to differences in repeat length, there is nothing common between these two tools despite the common CRISPR consensus motif.

Overall, diverse classes of repeats can be identified using a pipeline of tools. Differences in algorithmic nature reflect differences in repeat detection except for perfect microsatellites. Hence, the benchmarking of tools and development of the RDTcs server is essential and justified.

## 5. Conclusion

With the advancement of sequencing technologies, genome sequences have been continuously deposited in the repositories. Eventually, identification of repetitive sequences will be another follow-up task that can help to interpret the complex underlying mechanisms of disease, pathogenesis, and stress adaptation of biological systems. Understanding the diversity of repeat sequences and structures, various tools and algorithms have been developed to locate the repetitive patterns in the genome. Many of the users operate these tools without evaluating their efficiency and speed of repeat detection in diverse organisms. Furthermore, it is also very confusing to select a proper one from the huge pool of tools. Often, mistakes have been done in tool selection leading to limited repeat detection. To address this problem, a protocol has been standardized in the present work for the selection of specialized tools for each class of repeats. Repeat detection tools have been first classified and clustered into groups for the identification of tandem and interspersed repeats respectively. Then, for each group, rigorousness in repeat finding has been compared along with their execution time. 6 different biological systems comprising of 12 genomes have been opted to test the robustness of the tools. Also, the location and annotation of the repeats have been discussed. Among the perfect microsatellite identification tools, TROLL is selected for further use despite its limitations on input and outputs. This particular tool is preferred as it uses a dictionary of motifs leading to the correct estimation of the repeated space in a particular genome. Though TROLL is a little slower than SciRoko, Phobos, and PERF but can be compromised because of the benefit as mentioned above. In the present scenario, Phobos can be marked as the best tool for perfect minisatellites and imperfect repeat identification and has performed many times better than TRF both in repeat detection and execution time. Like Mreps, CRT detects many CRISPRs in prokaryotic genomes which introduce many false positives in the resulting repeat data. Hence, Piler-CR is selected for CRISPR detection. RepeatMasker has listed the maximum number of interspersed repeats in RepBase as compared to REPET hence preferred to use. The recommendation for tool selection for various types of repeats has been summarized in Table 4.

**Table 4:**
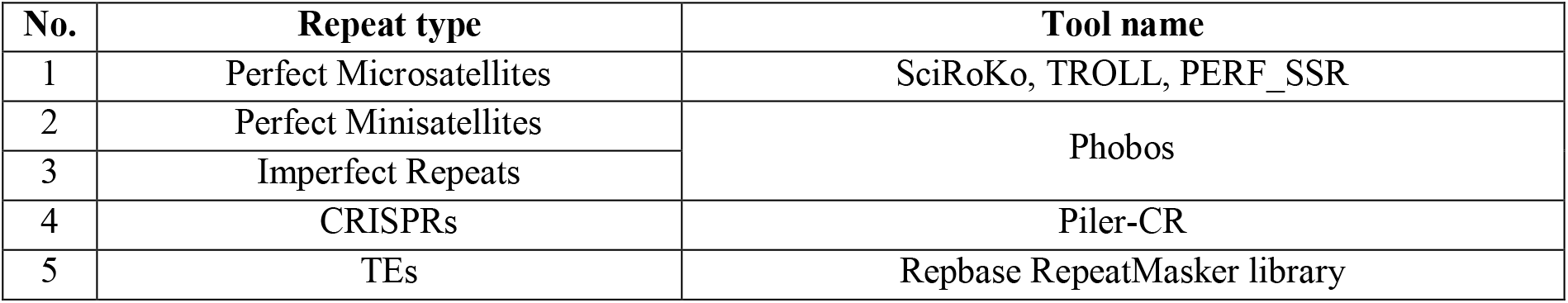
Recommendation for tool selection

In concluding remarks, it should be mentioned that equivalent parameter settings are required for proper comparison of tools. Survey, listing, and comparison of tools are essential for keeping up to date knowledge about the best tools. Moreover, the RDTcs server can be a very useful tool for designing markers for unknown disease-related genes creating pandemic like COVID-19.

## Supporting information

GDasRDTcsSupplementary

## Author’s Contribution

Conceptualization done by both, methodology, software, validation, formal analysis, investigation, resources, data curation, original draft preparation done by GD and review and editing, visualization, supervision, project administration is done by IG. All authors have read and agreed to the published version of the manuscript.

## Acknowledgments

GD is acknowledging the Council of Industrial Research (CSIR), Government of India for the senior research fellowship. We are thankful to Dr. Shailesh Kumar Panday and Dr. Pawan Kumar for checking the RDTcs web-server and their valuable suggestions.

## Supplementary Materials

GDasRDTcsSupplementaryAugust2020.pdf

## References

[1] G. Tóth, Z. Gáspári, J. Jurka, G. Toth, G. Tóth, Z. Gáspári, J. Jurka, Microsatellites in different eukaryotic genomes: survey and analysis., Genome Res. 10 (2000) 967–981. https://doi.org/10.1101/gr.10.7.967.

[2] S. Mehrotra, V. Goyal, Repetitive Sequences in Plant Nuclear DNA: Types, Distribution, Evolution and Function, Genomics, Proteomics Bioinforma. 12 (2014) 164–171. https://doi.org/10.1016/j.gpb.2014.07.003.

[3] C. Feschotte, N. Jiang, S.R. Wessler, Plant transposable elements: Where genetics meets genomics, Nat. Rev. Genet. 3 (2002) 329–341. https://doi.org/10.1038/nrg793.

[4] Z.N. Kronenberg, I.T. Fiddes, D. Gordon, S. Murali, S. Cantsilieris, O.S. Meyerson, J.G. Underwood, B.J. Nelson, M.J.P. Chaisson, M.L. Dougherty, K.M. Munson, A.R. Hastie, M. Diekhans, F. Hormozdiari, N. Lorusso, K. Hoekzema, R. Qiu, K. Clark, A. Raja, A.M.E. Welch, M. Sorensen, C. Baker, R.S. Fulton, J. Armstrong, T.A. Graves-Lindsay, A.M. Denli, E.R. Hoppe, P.H. Hsieh, C.M. Hill, A.W.C. Pang, J. Lee, E.T. Lam, S.K. Dutcher, F.H. Gage, W.C. Warren, J. Shendure, D. Haussler, V.A. Schneider, H. Cao, M. Ventura, R.K. Wilson, B. Paten, A. Pollen, E.E. Eichler, High-resolution comparative analysis of great ape genomes, Science (80-.). 360 (2018). https://doi.org/10.1126/science.aar6343.

[5] R. Gemayel, M.D. Vinces, M. Legendre, K.J. Verstrepen, Variable Tandem Repeats Accelerate Evolution of Coding and Regulatory Sequences, Annu. Rev. Genet. 44 (2010) 445–477. https://doi.org/10.1146/annurev-genet-072610-155046.

[6] R.K. Slotkin, R. Martienssen, Transposable elements and the epigenetic regulation of the genome., Nat. Rev. Genet. 8 (2007) 272–85. https://doi.org/10.1038/nrg2072.

[7] Y. Nakamura, K. Koyama, M. Matsushima, VNTR (variable number of tandem repeat) sequences as transcriptional, translational, or functional regulators., J. Hum. Genet. 43 (1998) 149–52. https://doi.org/10.1007/s100380050059.

[8] K. Usdin, The biological effects of simple tandem repeats: lessons from the repeat expansion diseases., Genome Res. 18 (2008) 1011–9. https://doi.org/10.1101/gr.070409.107.

[9] A.F. Saeed, R. Wang, S. Wang, Microsatellites in pursuit of microbial genome evolution, Front. Microbiol. 6 (2016) 1–15. https://doi.org/10.3389/fmicb.2015.01462.

[10] D. Waldron, Human evolution: Tandem repeats and divergent gene expression, Nat. Rev. Genet. 16 (2015) 7554. https://doi.org/10.1038/nrg4040.

[11] G. Das, S. Das, S. Dutta, I. Ghosh, In silico identification and characterization of stress and virulence associated repeats in Salmonella, Genomics. 110 (2018) 23–34. https://doi.org/10.1016/j.ygeno.2017.08.002.

[12] A. Merkel, N. Gemmell, Detecting short tandem repeats from genome data: Opening the software black box, Brief. Bioinform. 9 (2008) 355–366. https://doi.org/10.1093/bib/bbn028.

[13] K.G. Lim, C.K. Kwoh, L.Y. Hsu, A. Wirawan, Review of tandem repeat search tools: A systematic approach to evaluating algorithmic performance, Brief. Bioinform. 14 (2013) 67–81. https://doi.org/10.1093/bib/bbs023.

[14] S. Saha, S. Bridges, Z. V. Magbanua, D.G. Peterson, Empirical comparison of ab initio repeat finding programs, Nucleic Acids Res. 36 (2008) 2284–2294. https://doi.org/10.1093/nar/gkn064.

[15] P.C. Sharma, A. Grover, G. Kahl, Mining microsatellites in eukaryotic genomes, Trends Biotechnol. 25 (2007) 490–498. https://doi.org/10.1016/j.tibtech.2007.07.013.

[16] E. Lerat, Identifying repeats and transposable elements in sequenced genomes: how to find your way through the dense forest of programs., Heredity (Edinb). 104 (2010) 520–533. https://doi.org/10.1038/hdy.2009.165.

[17] S. Saha, S. Bridges, Z. V. Magbanua, D.G. Peterson, Computational Approaches and Tools Used in Identification of Dispersed Repetitive DNA Sequences, Trop. Plant Biol. 1 (2008) 85–96. https://doi.org/10.1007/s12042-007-9007-5.

[18] P. Goerner-Potvin, G. Bourque, Computational tools to unmask transposable elements, Nat. Rev. Genet. 19 (2018) 688–704. https://doi.org/10.1038/s41576-018-0050-x.

[19] L. Rishishwar, L. Mariño-Ramírez, I.K. Jordan, Benchmarking computational tools for polymorphic transposable element detection, Brief. Bioinform. 18 (2017) 908–918. https://doi.org/10.1093/bib/bbw072.

[20] V.J. Henry, A.E. Bandrowski, A.S. Pepin, B.J. Gonzalez, A. Desfeux, OMICtools: an informative directory for multi-omic data analysis, Database (Oxford). 2014 (2014) 1–5. https://doi.org/10.1093/database/bau069.

[21] N.A. O’Leary, M.W. Wright, J.R. Brister, S. Ciufo, D. Haddad, R. McVeigh, B. Rajput, B. Robbertse, B. Smith-White, D. Ako-Adjei, A. Astashyn, A. Badretdin, Y. Bao, O. Blinkova, V. Brover, V. Chetvernin, J. Choi, E. Cox, O. Ermolaeva, C.M. Farrell, T. Goldfarb, T. Gupta, D. Haft, E. Hatcher, W. Hlavina, V.S. Joardar, V.K. Kodali, W. Li, D. Maglott, P. Masterson, K.M. McGarvey, M.R. Murphy, K. O’Neill, S. Pujar, S.H. Rangwala, D. Rausch, L.D. Riddick, C. Schoch, A. Shkeda, S.S. Storz, H. Sun, F. Thibaud-Nissen, I. Tolstoy, R.E. Tully, A.R. Vatsan, C. Wallin, D. Webb, W. Wu, M.J. Landrum, A. Kimchi, T. Tatusova, M. DiCuccio, P. Kitts, T.D. Murphy, K.D. Pruitt, Reference sequence (RefSeq) database at NCBI: Current status, taxonomic expansion, and functional annotation, Nucleic Acids Res. 44 (2016) D733–D745. https://doi.org/10.1093/nar/gkv1189.

[22] G. Benson, Tandem repeats finder: A program to analyze DNA sequences, Nucleic Acids Res. 27 (1999) 573–580. https://doi.org/10.1093/nar/27.2.573.

[23] A.T. Castelo, W. Martins, G.R. Gao, TROLL--Tandem Repeat Occurrence Locator, Bioinformatics. 18 (2002) 634–636. https://doi.org/10.1093/bioinformatics/18.4.634.

[24] Y. Gelfand, A. Rodriguez, G. Benson, TRDB - The Tandem Repeats Database, Nucleic Acids Res. 35 (2007). https://doi.org/10.1093/nar/gkl1013.

[25] T. Thiel, W. Michalek, R. Varshney, A. Graner, Exploiting EST databases for the development and characterization of gene-derived SSR-markers in barley (Hordeum vulgare L.), Theor. Appl. Genet. 106 (2003) 411–422. https://doi.org/10.1007/s00122-002-1031-0.

[26] S. Beier, T. Thiel, T. Münch, U. Scholz, M. Mascher, MISA-web: a web server for microsatellite prediction, Bioinformatics. 33 (2017) 2583–2585. https://doi.org/10.1093/bioinformatics/btx198.

[27] A. Merkel, N.J. Gemmell, A. Merkel, N.J. Gemmell, Detecting Microsatellites in Genome Data: Variance in Definitions and Bioinformatic Approaches Cause Systematic Bias, Evol. Bioinforma. (2008). https://doi.org/10.4137/EBO.S420.

[28] C. Mayer, F. Leese, R. Tollrian, Genome-wide analysis of tandem repeats in Daphnia pulex - a comparative approach, BMC Genomics. 11 (2010). https://doi.org/10.1186/1471-2164-11-277.

[29] S. Temnykh, G. DeClerck, A. Lukashova, L. Lipovich, S. Cartinhour, S. McCouch, Computational and experimental analysis of microsatellites in rice (Oryza sativa L.): Frequency, length variation, transposon associations, and genetic marker potential, Genome Res. 11 (2001) 1441–1452. https://doi.org/10.1101/gr.184001.

[30] R. Kofler, C. Schlötterer, T. Lelley, SciRoKo: A new tool for whole genome microsatellite search and investigation, Bioinformatics. 23 (2007) 1683–1685. https://doi.org/10.1093/bioinformatics/btm157.

[31] A.K. Avvaru, D.T. Sowpati, R.K. Mishra, PERF: An exhaustive algorithm for ultra-fast and efficient identification of microsatellites from large DNA sequences, Bioinformatics. (2018). https://doi.org/10.1093/bioinformatics/btx721.

[32] S. Beier, T. Thiel, T. Münch, U. Scholz, M. Mascher, MISA-web: a web server for microsatellite prediction, Bioinformatics. 33 (2017) 2583–2585. https://doi.org/10.1093/bioinformatics/btx198.

[33] G. Achaz, E.P.C. Rocha, P. Netter, E. Coissac, Origin and fate of repeats in bacteria., Nucleic Acids Res. 30 (2002) 2987–2994. https://doi.org/10.1093/nar/gkf391.

[34] R. Kolpakov, G. Bana, G. Kucherov, mreps: Efficient and flexible detection of tandem repeats in DNA, Nucleic Acids Res. 31 (2003) 3672–3678. https://doi.org/10.1093/nar/gkg617.

[35] C. Bland, T.L. Ramsey, F. Sabree, M. Lowe, K. Brown, N.C. Kyrpides, P. Hugenholtz, CRISPR Recognition Tool (CRT): A tool for automatic detection of clustered regularly interspaced palindromic repeats, BMC Bioinformatics. 8 (2007). https://doi.org/10.1186/1471-2105-8-209.

[36] A. Biswas, R.H.J. Staals, S.E. Morales, P.C. Fineran, C.M. Brown, CRISPRDetect: A flexible algorithm to define CRISPR arrays, BMC Genomics. 17 (2016) 356. https://doi.org/10.1186/s12864-016-2627-0.

[37] R.C. Edgar, PILER-CR: fast and accurate identification of CRISPR repeats., BMC Bioinformatics. 8 (2007) 1–6. https://doi.org/10.1186/1471-2105-8-18.

[38] L. Rishishwar, L. Mariño-Ramírez, I.K. Jordan, Benchmarking computational tools for polymorphic transposable element detection, Brief. Bioinform. (2016) bbw072. https://doi.org/10.1093/bib/bbw072.

[39] W. Bao, K.K. Kojima, O. Kohany, Repbase Update, a database of repetitive elements in eukaryotic genomes, Mob. DNA. 6 (2015) 11. https://doi.org/10.1186/s13100-015-0041-9.

[40] M. Tarailo-Graovac, N. Chen, Using RepeatMasker to identify repetitive elements in genomic sequences, Curr. Protoc. Bioinforma. (2009). https://doi.org/10.1002/0471250953.bi0410s25.

[41] T. Flutre, E. Duprat, C. Feuillet, H. Quesneville, Considering transposable element diversification in de novo annotation approaches, PLoS One. (2011). https://doi.org/10.1371/journal.pone.0016526.

[42] J.A. Shapiro, R. von Sternberg, Why repetitive DNA is essential to genome function., Biol. Rev. Camb. Philos. Soc. 80 (2005) 227–50. https://doi.org/10.1017/S1464793104006657.

[43] A.R. Quinlan, I.M. Hall, BEDTools: A flexible suite of utilities for comparing genomic features, Bioinformatics. 26 (2010) 841–842. https://doi.org/10.1093/bioinformatics/btq033.

[44] J.R. Conway, A. Lex, N. Gehlenborg, UpSetR: An R package for the visualization of intersecting sets and their properties, Bioinformatics. 33 (2017) 2938–2940. https://doi.org/10.1093/bioinformatics/btx364.

[45] V.J. Henry, A.E. Bandrowski, A.S. Pepin, B.J. Gonzalez, A. Desfeux, OMICtools: an informative directory for multi-omic data analysis, Database (Oxford). 2014 (2014). https://doi.org/10.1093/database/bau069.

