## Supplementary material for "Benchmarking tools for DNA repeat identification in diverse genomes": GDasRDTcsSupplementary

### Supplementary Table and Figures


| | Page No. |
|---|---|
| Table | 2 |
| Figures | 3-20 |

Table T1: Comparing outputs of the tools installed in local computer with published results

| No. | Tool | Input Sequences (Genbank/RefSeq Accession) | Published results | Current result | PMID |
|---|---|---|---|---|---|
| 1 | MISA | AC256511.1, AC269605.1, AC265197.1, AC263353.1, AC264961.1, AC266636.1, AC261250.1, AC267178.1, AC259365.1, AC257258.1 | 6022 | 6022 | 28398459 |
| 2 | TROLL | | 15091 | 15091 | |
| 3 | SciRoko | | 6021 | 6021 | |
| 4 | SSRIT | NC_000001.11 | 304764 | 304764 | 29121165 |
| 5 | PERF | | 350932 | 350932 | |
| 6 | Mreps | | 172449 | 172449 | |
| 7 | TRF | KU179220.1 | 28 | 28 | 28326093 |
| 8 | Phobos | | 419 | 419 | |
| 9 | CRISPR-Detect | NC_003198.1 | No. of arrays: 1<br>Repeat no.: 8 | No. of arrays: 1<br>Repeat no.: 8 | 27184979 |
| 10 | PILER-CR | | No. of arrays: 1<br>Repeat no.: 6 | No. of arrays: 1<br>Repeat no.: 6 | |
| 11 | CRT | | No. of arrays: 1<br>Repeat no.: 7 | No. of arrays: 1<br>Repeat no.: 7 | |

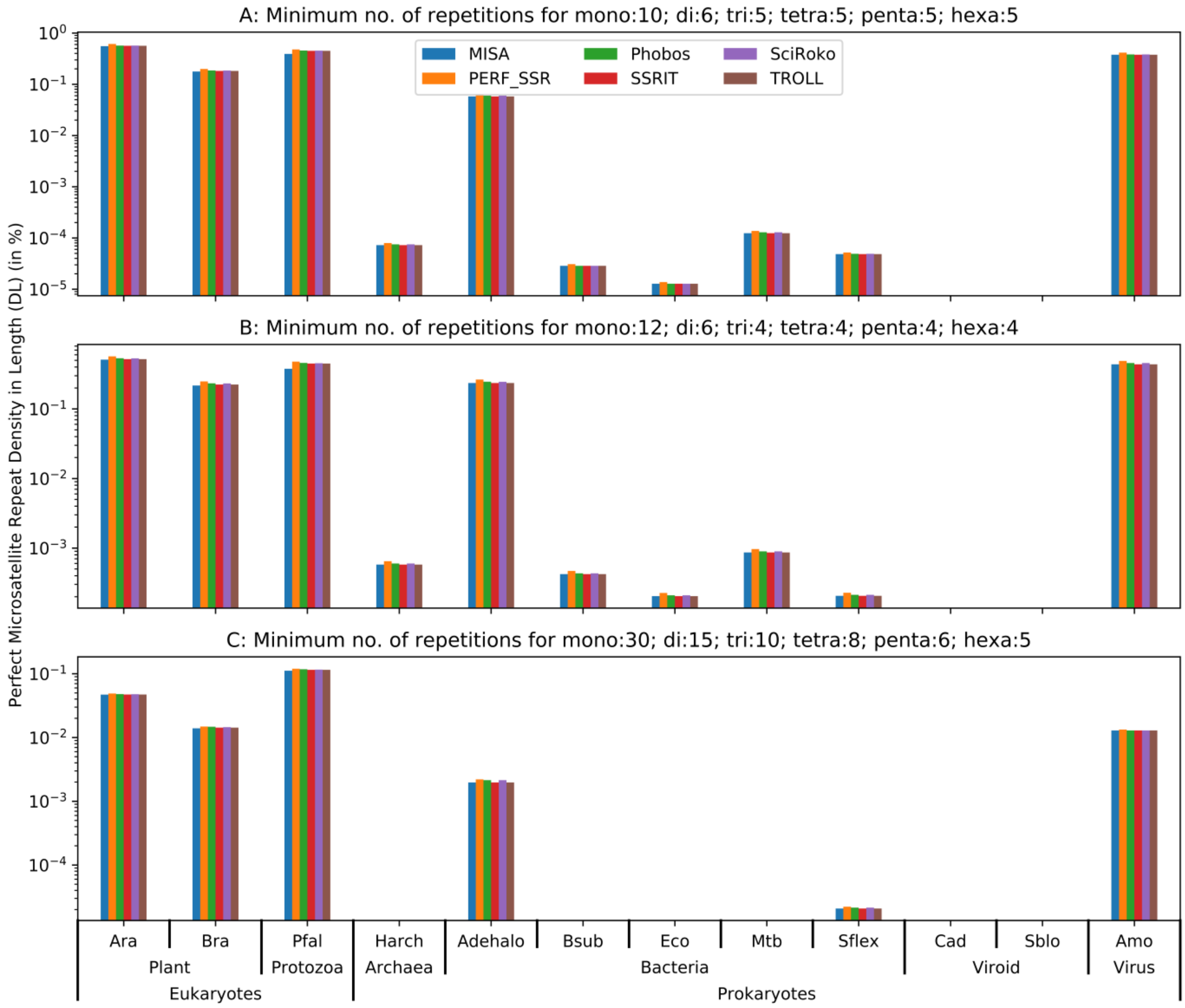


Figure F1: Comparison of repeat density in length parameter ($D_L$) of perfect microsatellite detection tools using a diverse set of genomes and 3 distinct set of values for repeat no. parameter. Y-axes are in log scale.

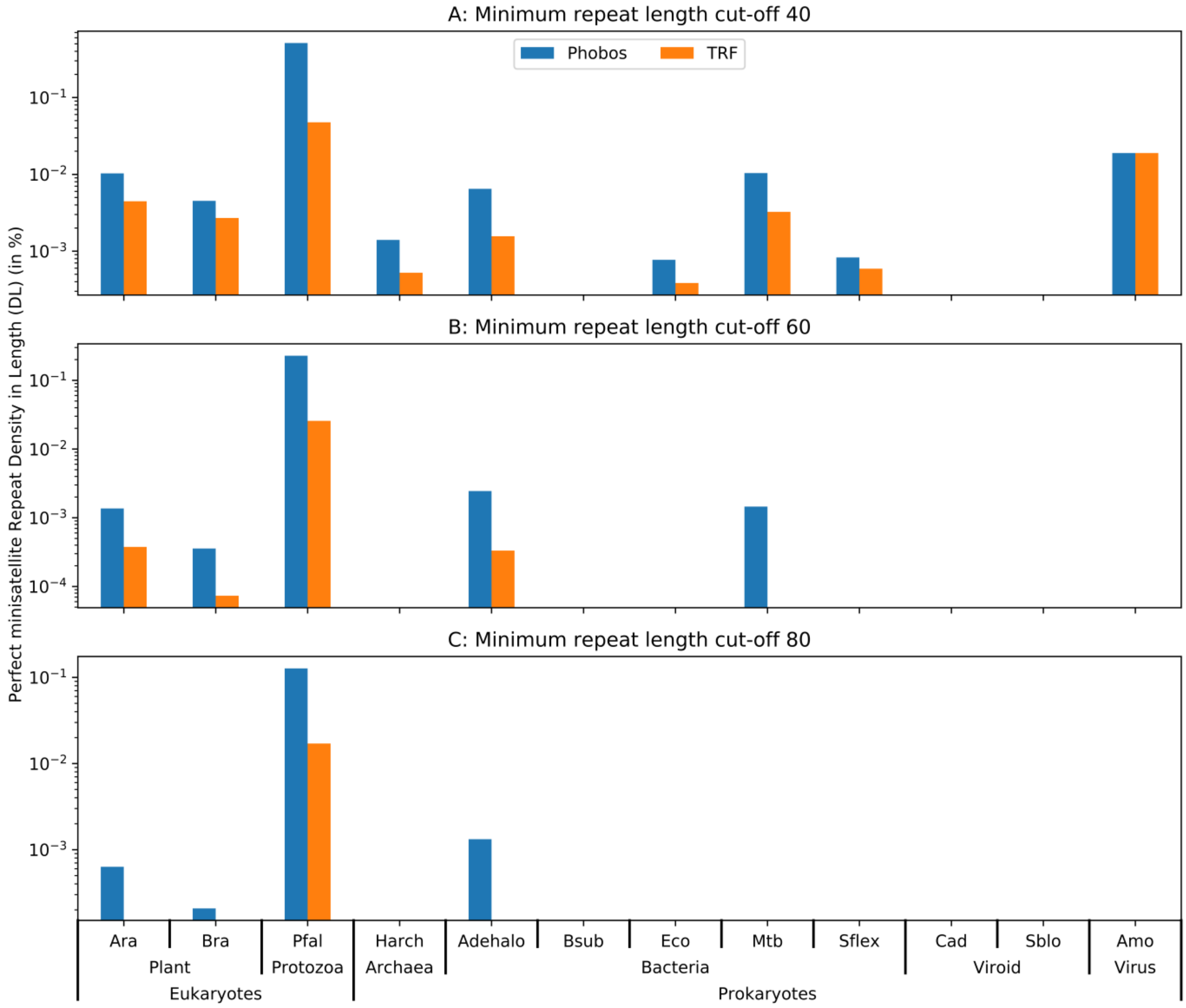


Figure F2: Comparison of repeat density in length parameter ($D_L$) of perfect minisatellite detection tools using a diverse set of genomes and 3 distinct set of values for repeat no. parameter. Y-axes are in log scale.

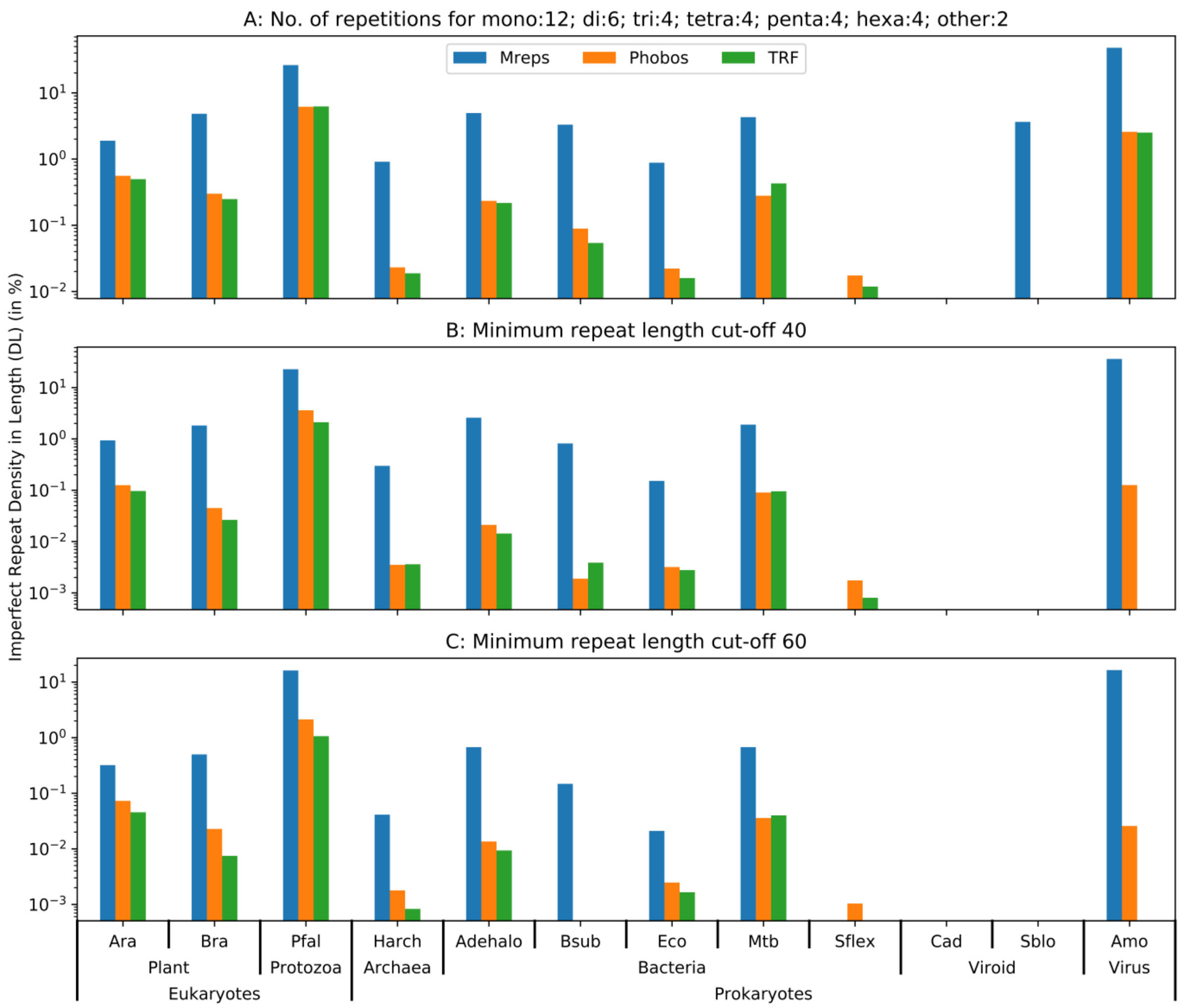


Figure F3: Comparison of repeat density in length parameter ($D_L$) of imperfect tandem repeat detection tools using a diverse set of genomes and 3 distinct set of values for repeat no. parameter. Y-axes are in log scale.

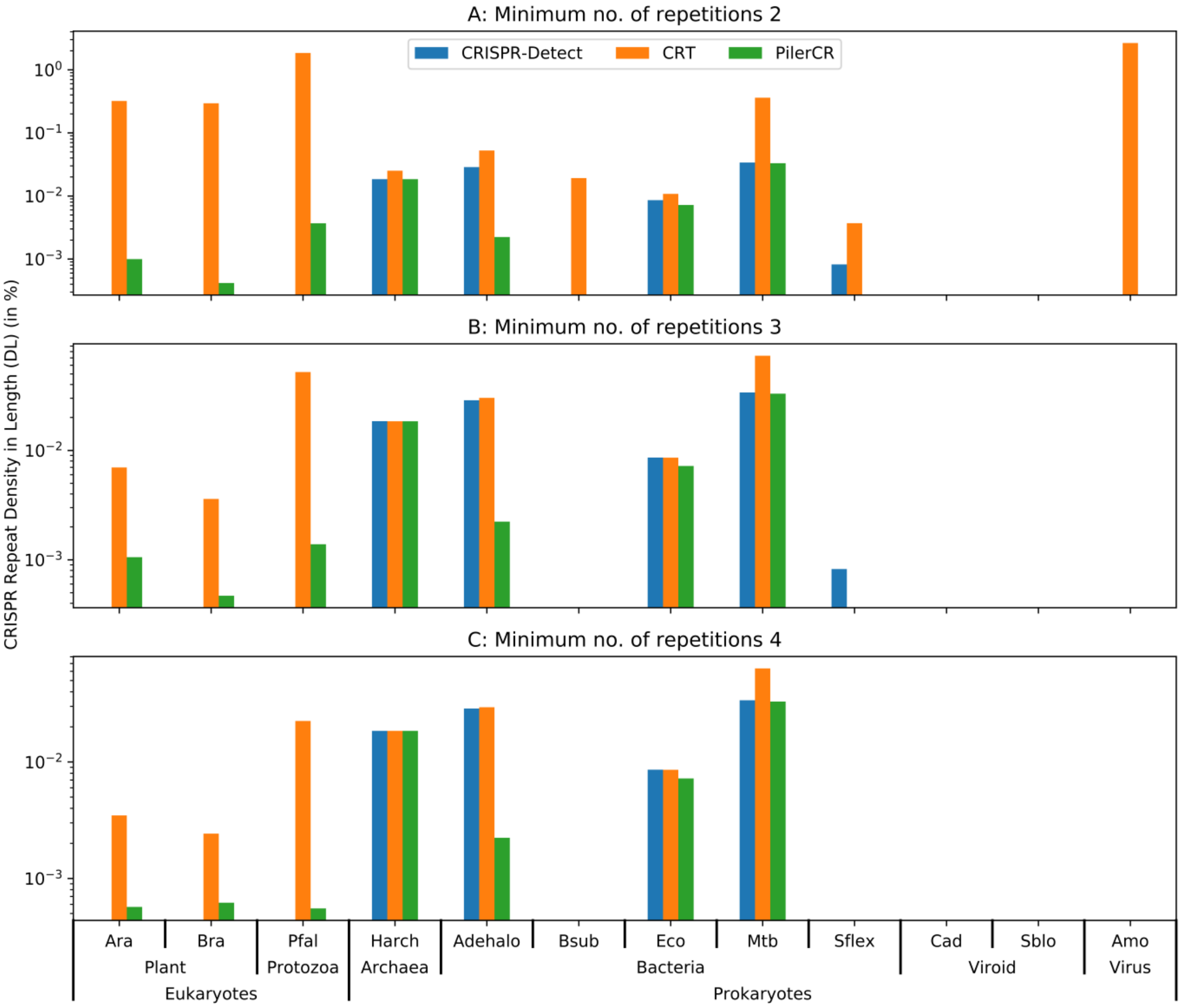


Figure F4: Comparison of repeat density in length parameter ($D_L$) of CRISPR detection tools using a diverse set of genomes and 3 distinct set of values for repeat no. parameter. Y-axes are in log scale.

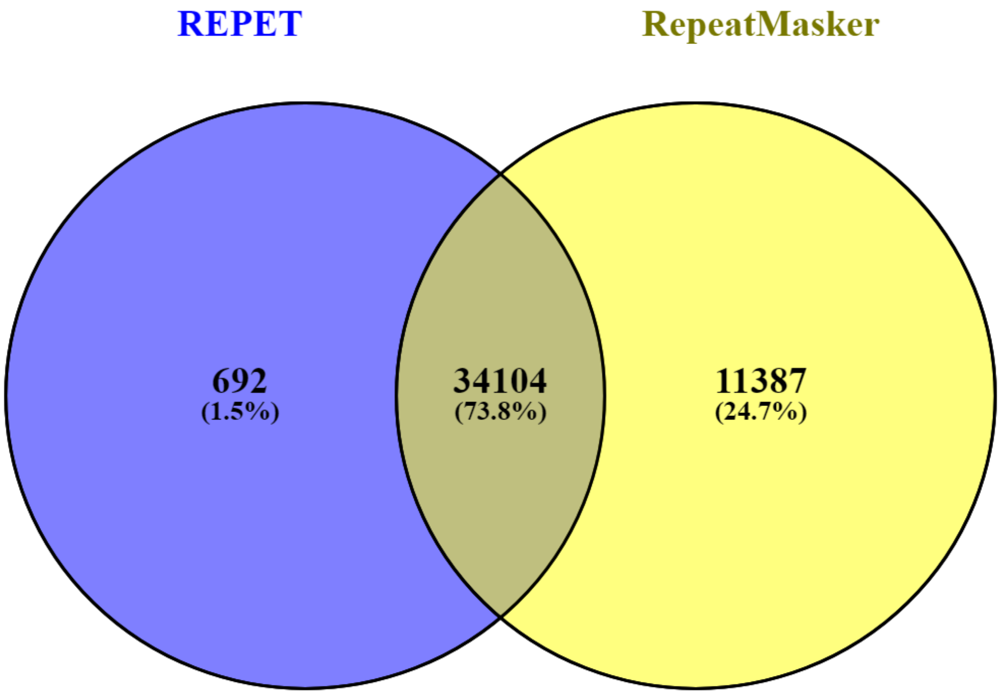


Figure F5: Venn diagram of interspersed repeats extracted from the REPET (blue) and RepeatMasker (yellow) libraries present in Repbase database. Unique and common entries have been represented in numbers and percentages in brackets.

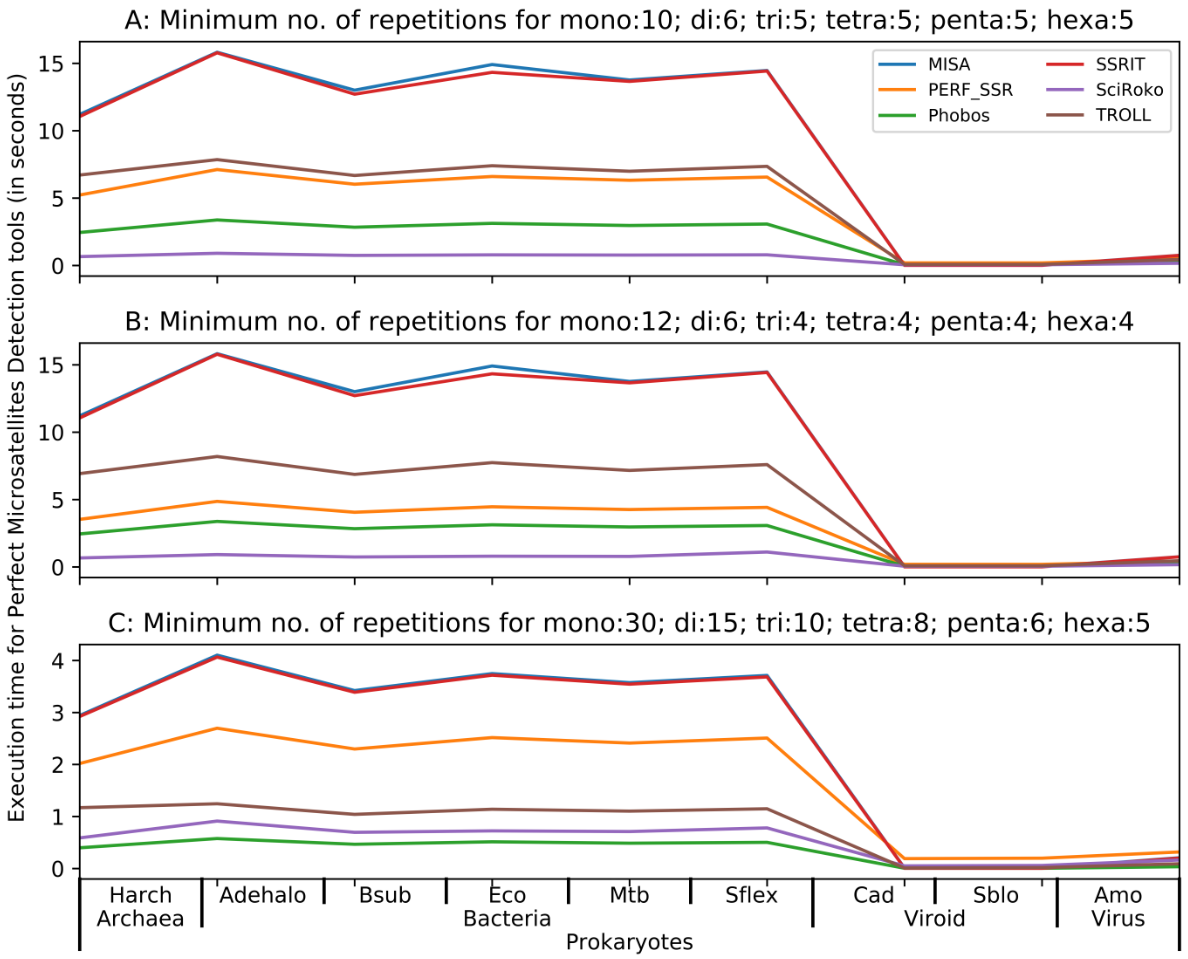


Figure F6: Comparison of tool's execution time for perfect microsatellite extraction in prokaryotes using 3 different set of values of program parameters. Y-axis is in seconds.

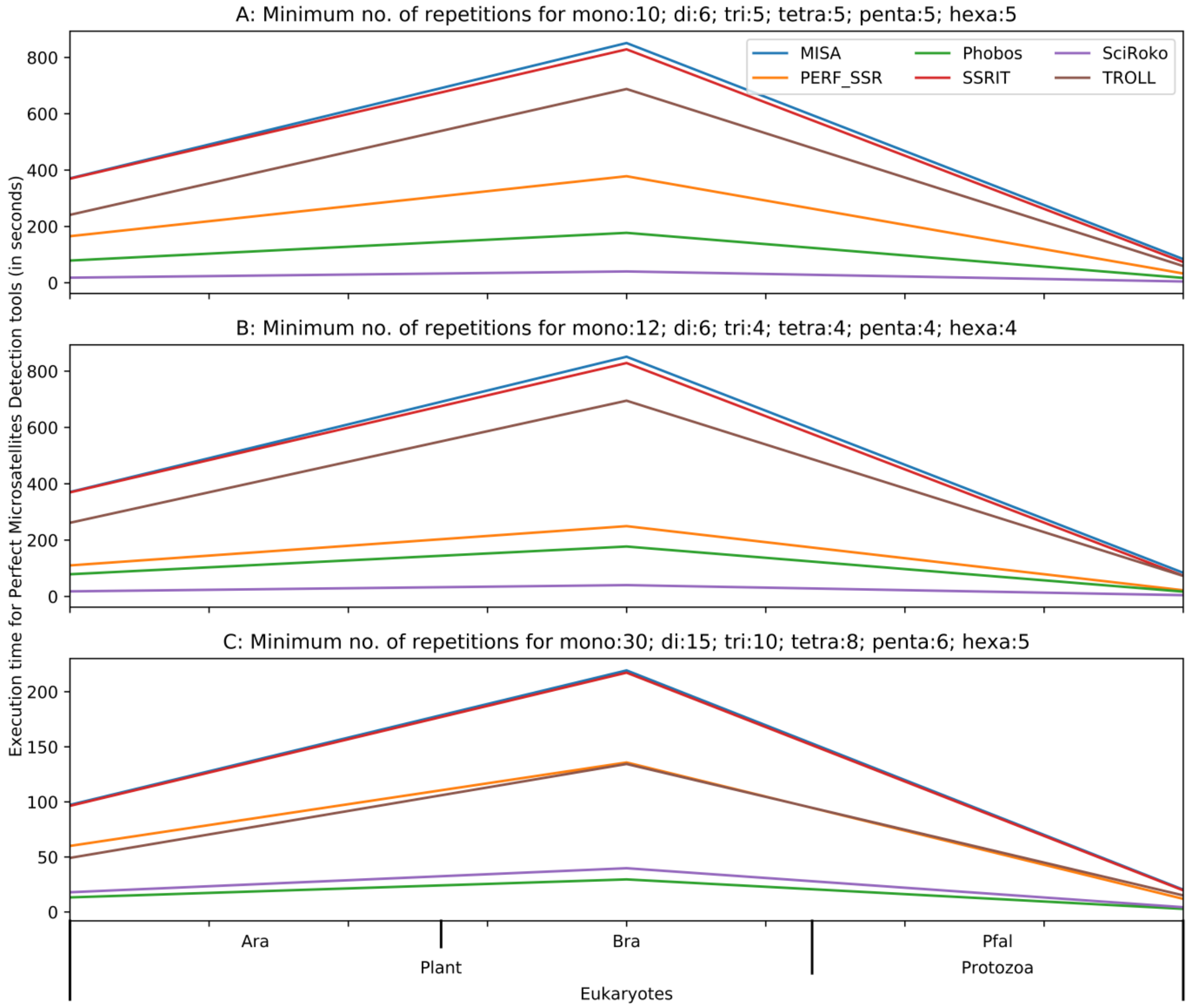


Figure F7: Comparison of tool's execution time for perfect microsatellite extraction in eukaryotes using 3 different set of values of program parameters. Y-axis is in seconds.

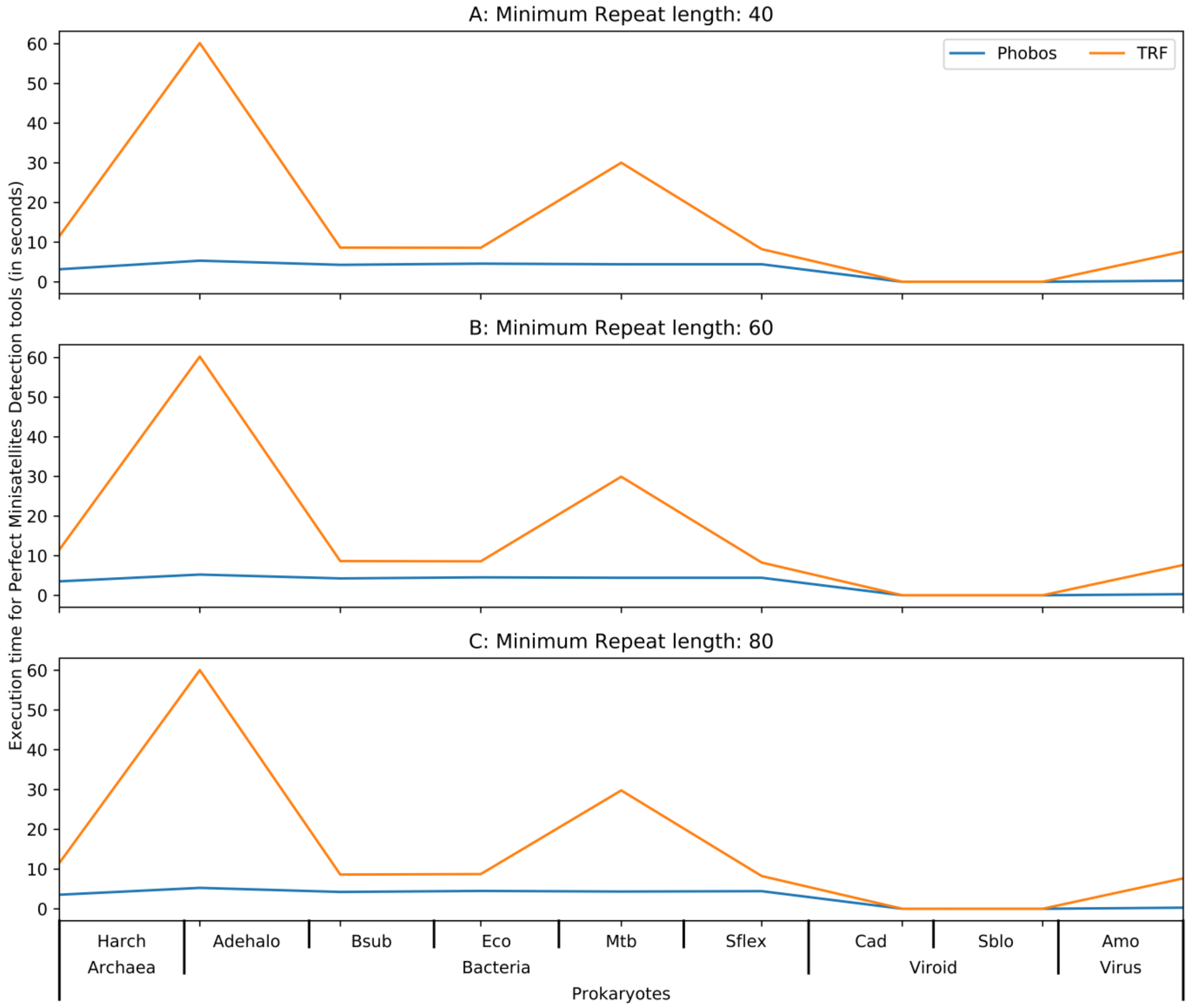


Figure F8: Comparison of tool's execution time for perfect minisatellite extraction in prokaryotes using 3 different set of values of program parameters. Y-axis is in seconds.

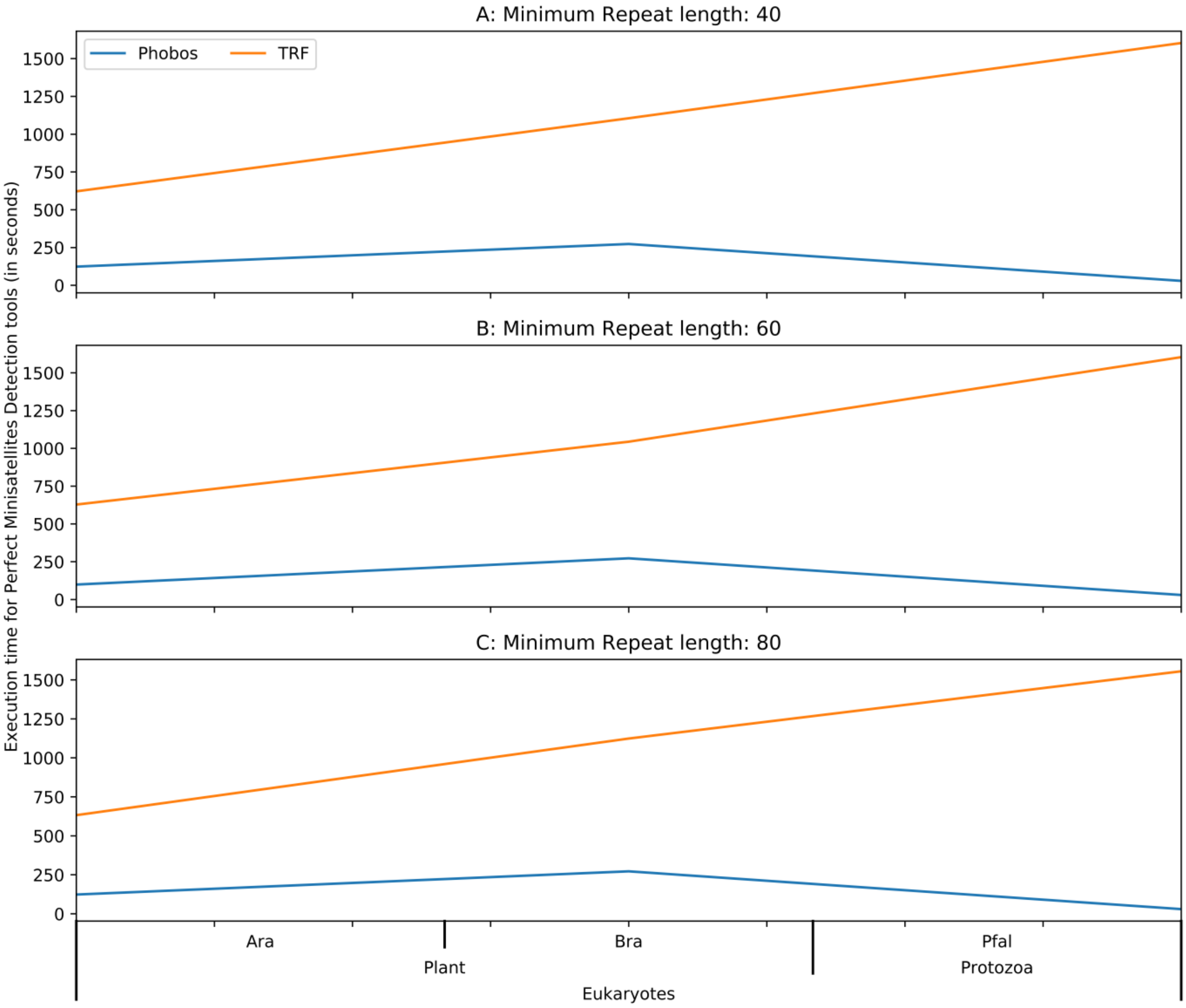


Figure F9: Comparison of tool's execution time for perfect minisatellite extraction in eukaryotes using 3 different set of values of program parameters. Y-axis is in seconds.

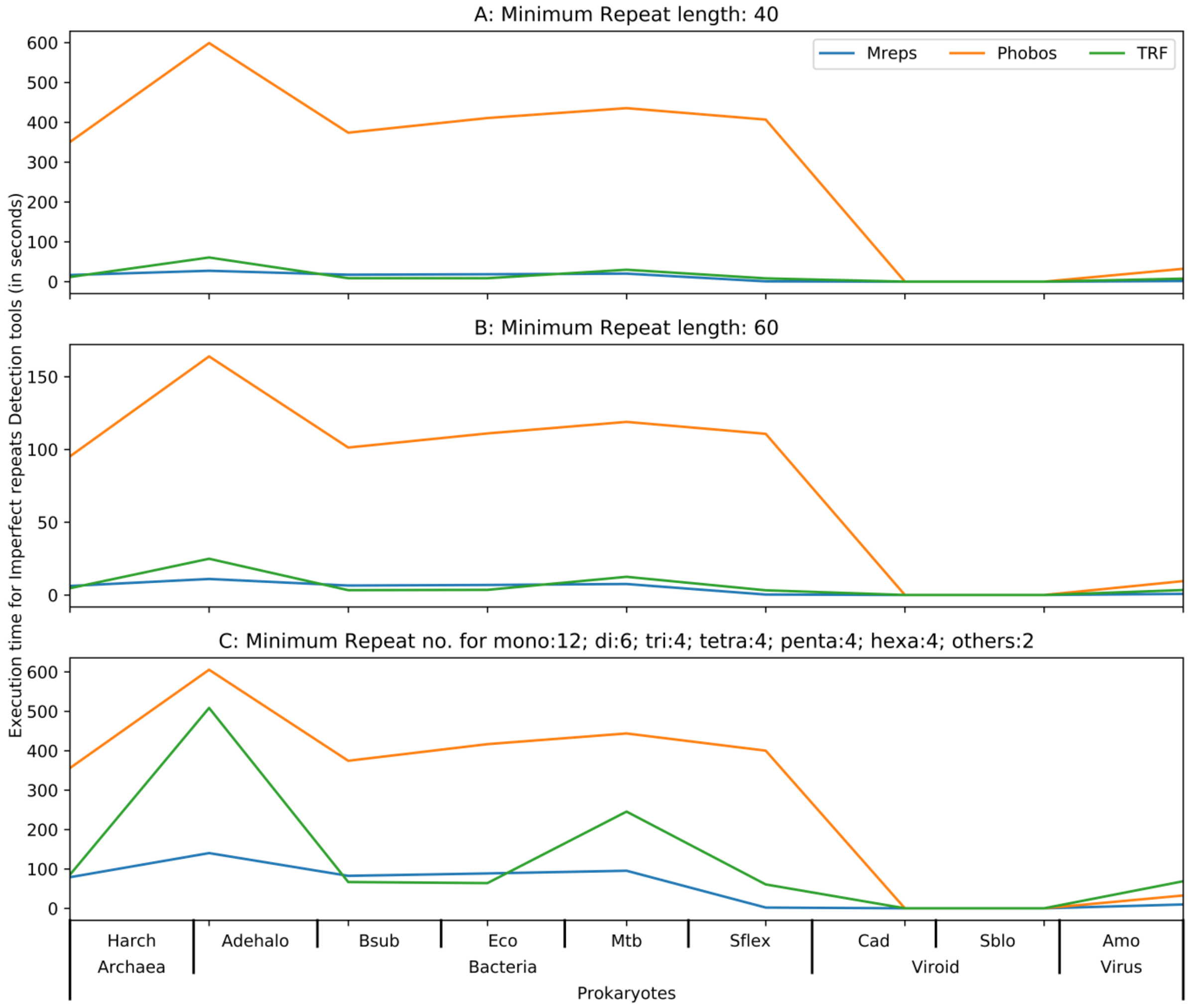


Figure F10**:** Comparison of tool's execution time for imperfect repeats extraction in prokaryotes using 3 different set of values of program parameters. Y-axis is in seconds.

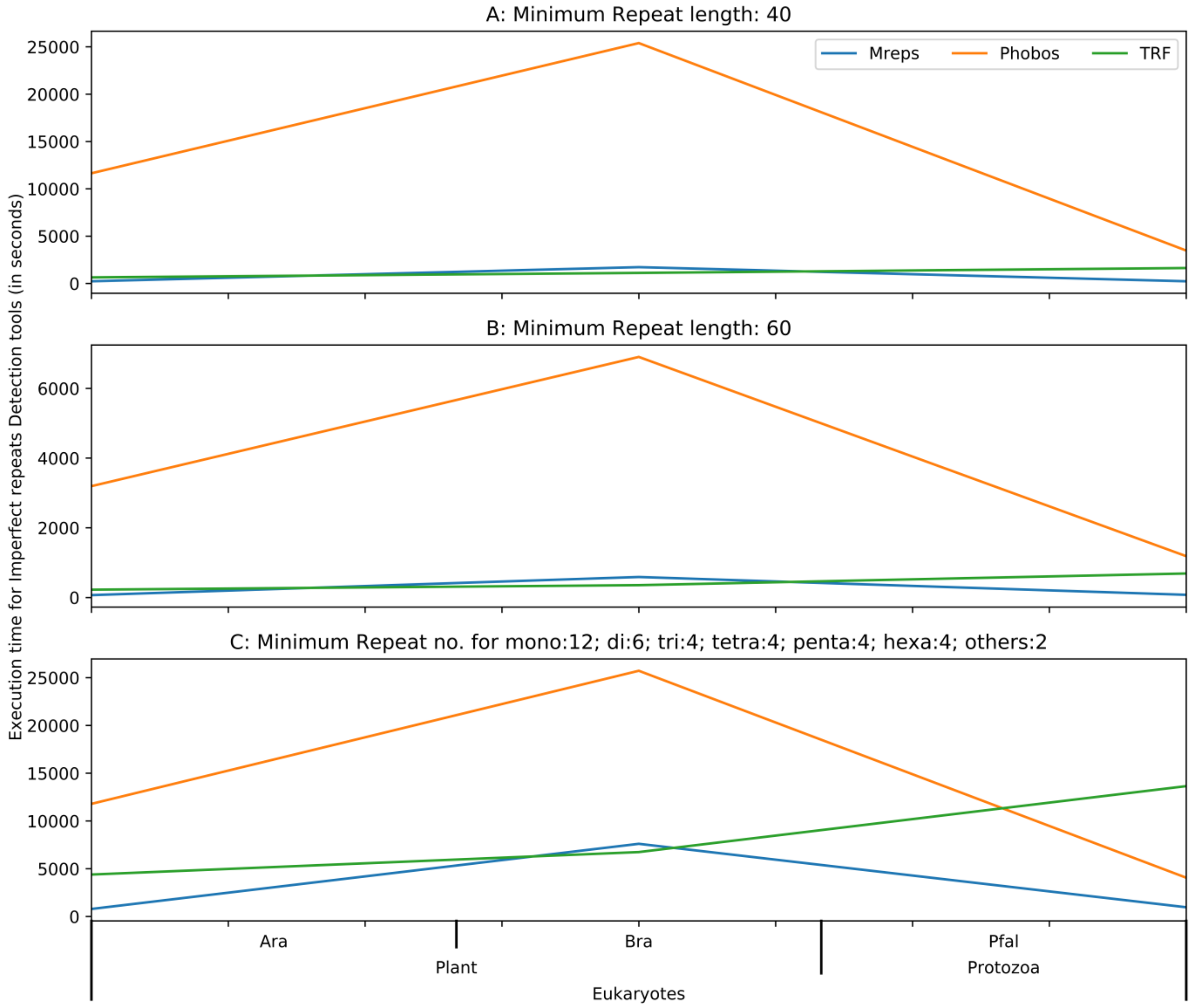


Figure F11: Comparison of tool's execution time for imperfect repeats extraction in eukaryotes using 3 different set of values of program parameters. Y-axis is in seconds.

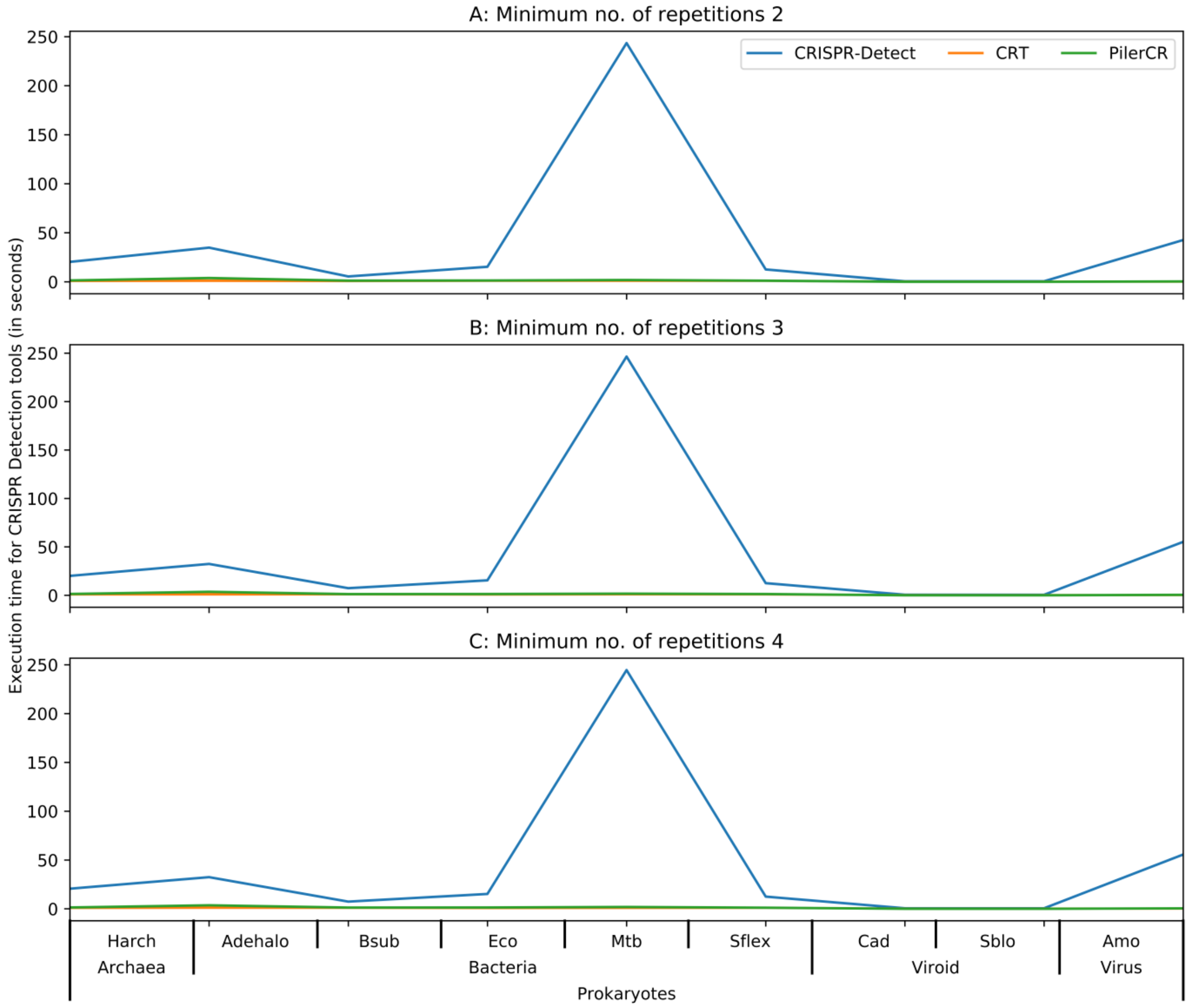


Figure F12: Comparison of tool's execution time for CRISPRs extraction in prokaryotes using 3 different set of values of program parameters. Y-axis is in seconds.

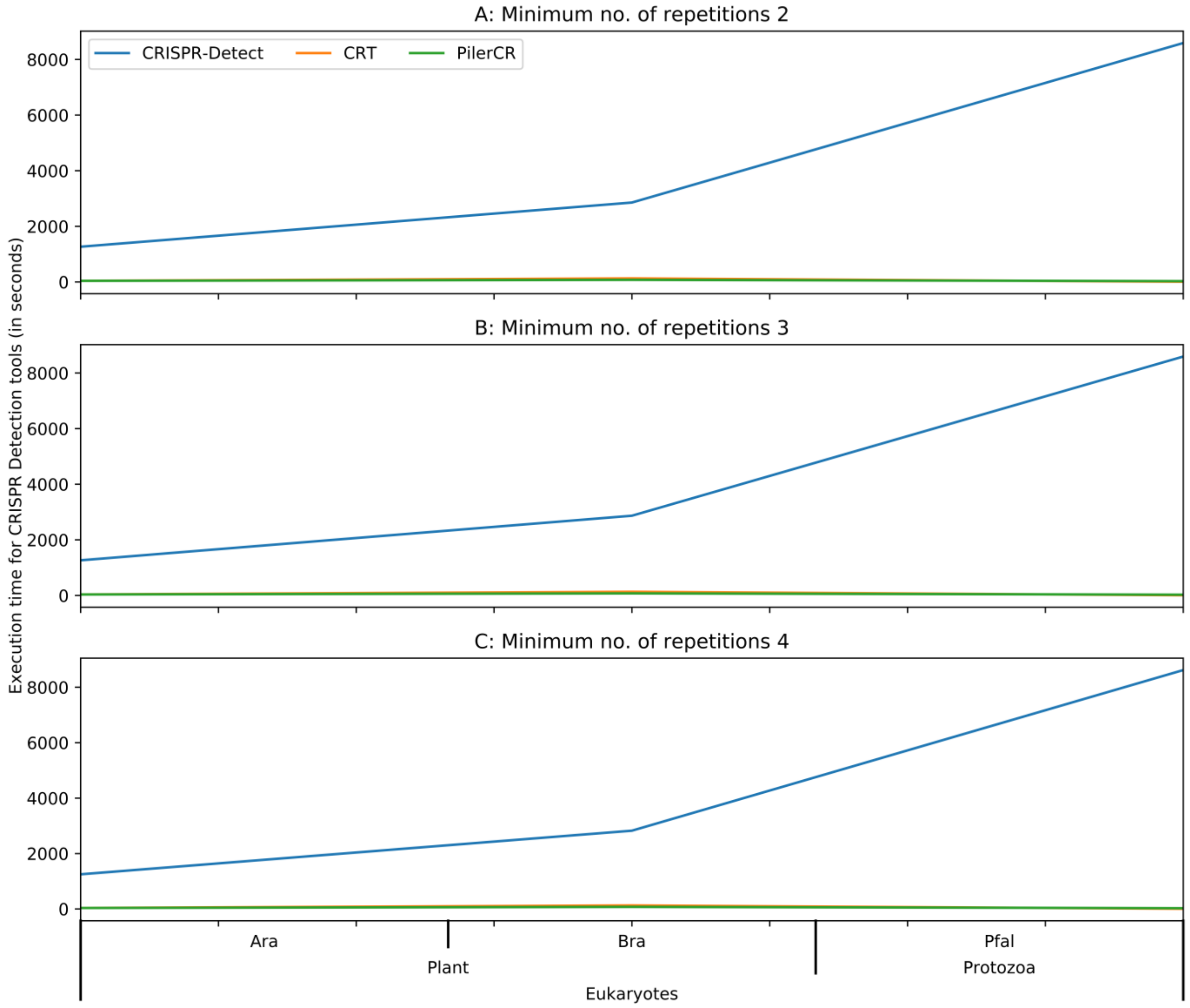


Figure F13: Comparison of tool’s execution time for CRISPRS extraction in eukaryotes using 3 different set of values of program parameters. Y-axis is in seconds.

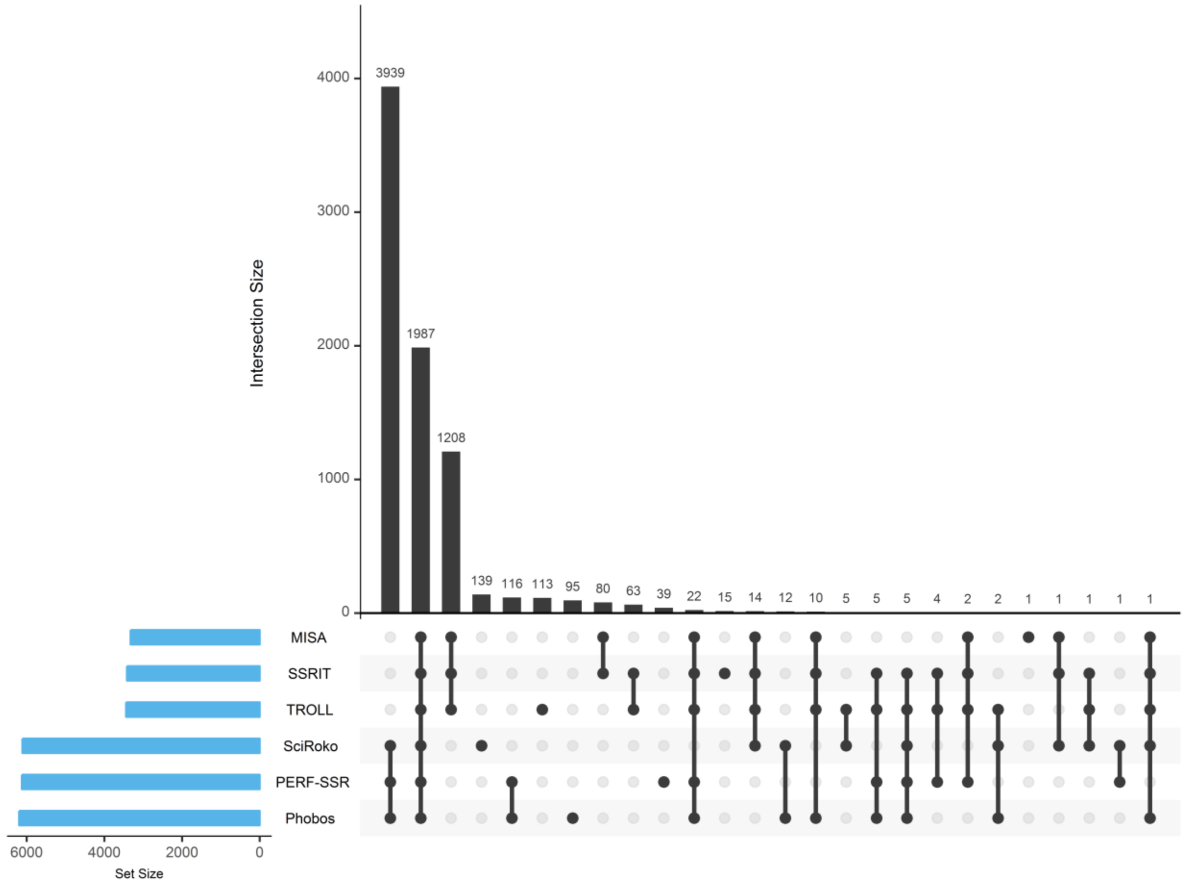


Figure F14: Analysis of cumulative perfect microsatellite repeats extracted by all tools using aforementioned parameter values. Motif and repeat length have been used together while counting. Zero intersections have not shown in the plot.

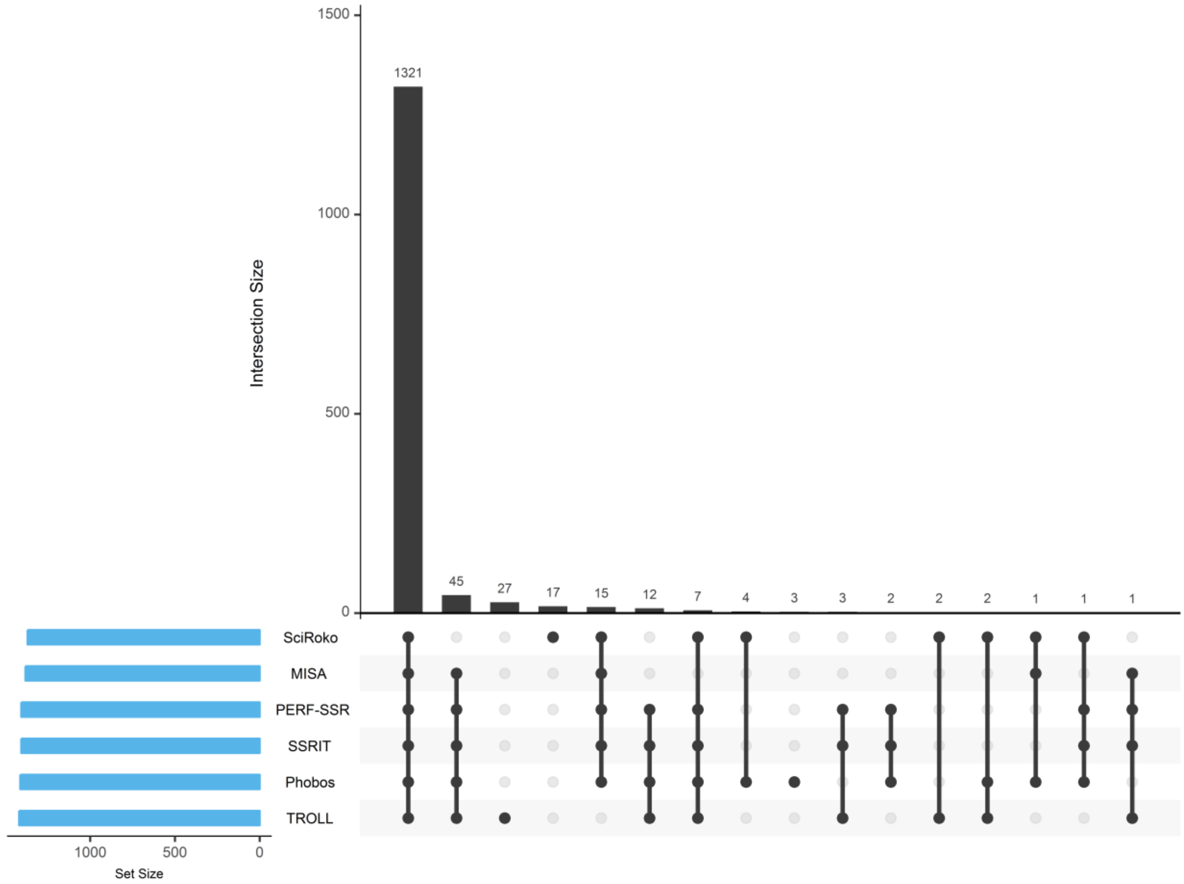


Figure F15: Analysis of cumulative perfect microsatellite repeating motifs extracted by all tools using aforementioned parameter values. Only motif information has been used while counting. Zero intersections have not shown in the plot.

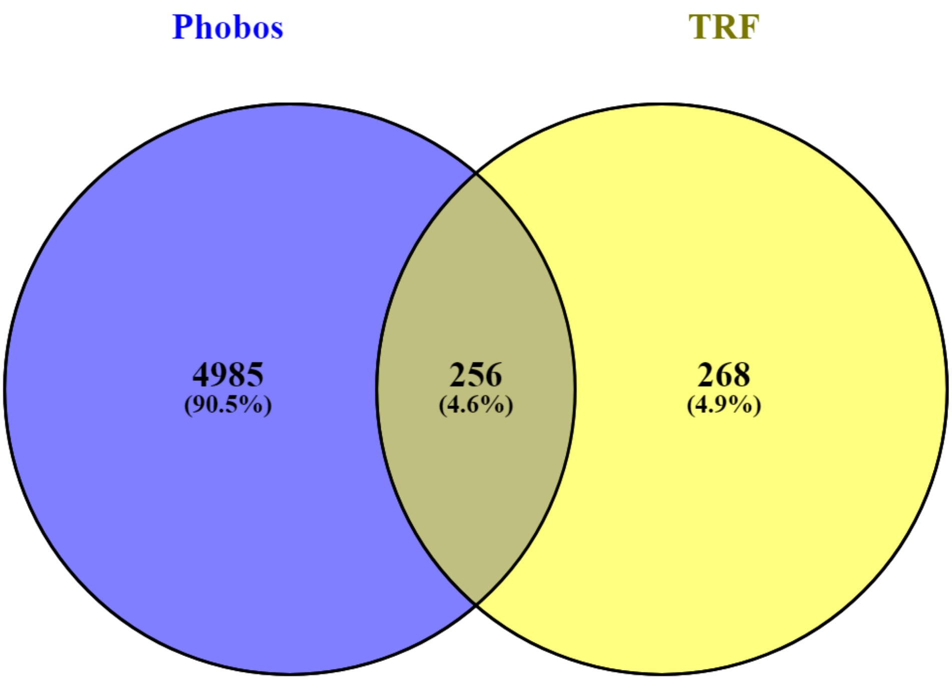


Figure F16: Venn diagram of all perfect minisatellites repeats extracted by tools Phobos (blue) and TRF (yellow). Unique and common entries have been represented in numbers and percentages in brackets. Motif and repeat length have been used together while counting.

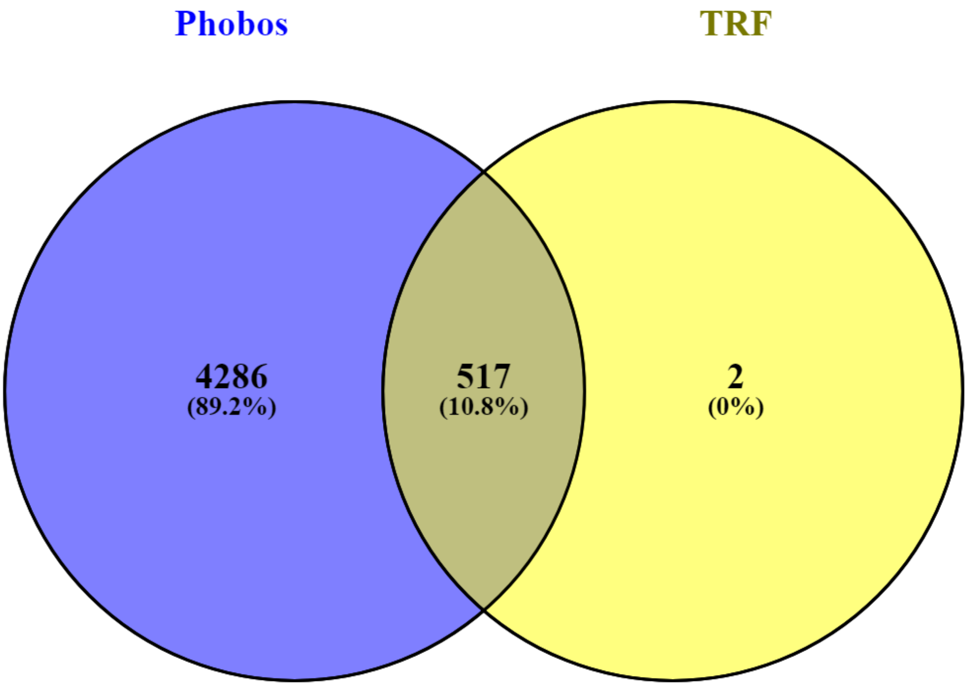


Figure F17: Venn diagram of all perfect minisatellites repeats extracted by tools Phobos (blue) and TRF (yellow). Unique and common entries have been represented in numbers and percentages in brackets. Motif information has been used only while counting. Only 1st decimal places have been considered that is why 0% is showing. Indeed it's a very small percentage.

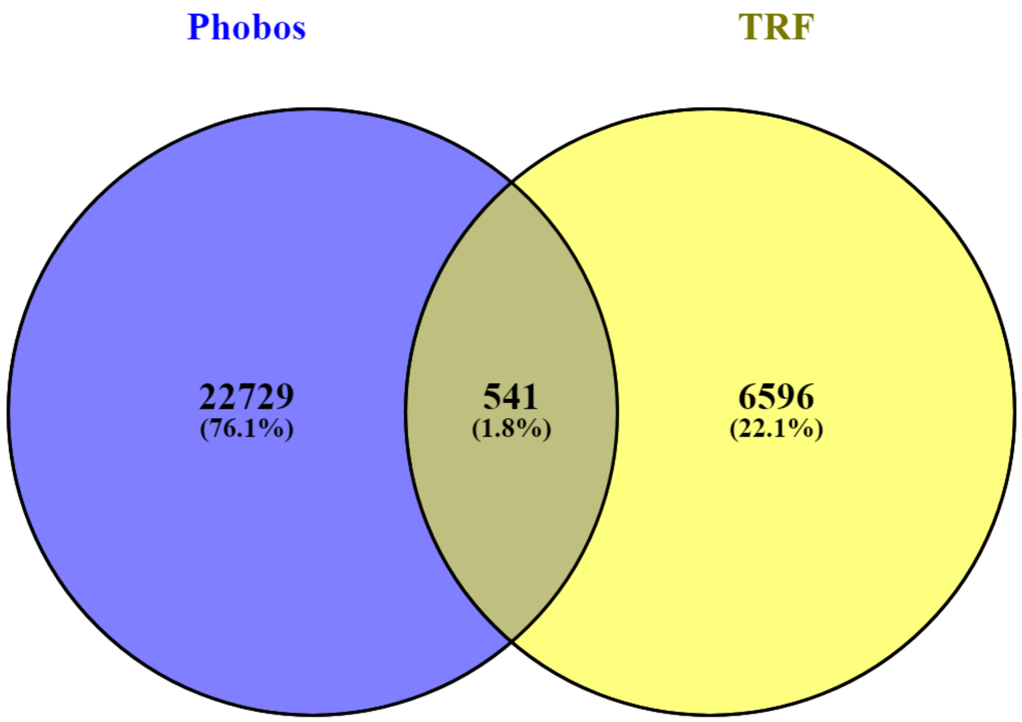


Figure F18: Venn diagram of all imperfect repeats extracted by tools Phobos (blue) and TRF (yellow). Mreps do not report any motif information hence not considered. Unique and common entries have been represented in numbers and percentages in brackets. Motif and repeat length have been used together while counting.

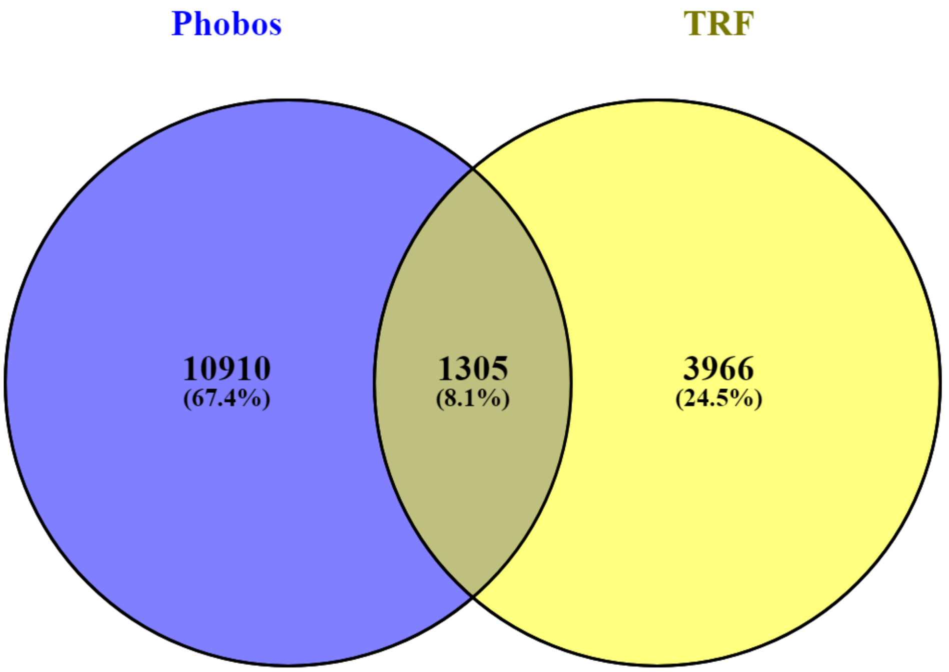


Figure F19: Venn diagram of all imperfect repeats extracted by tools Phobos (blue) and TRF (yellow). Mreps do not report any motif information hence not considered. Unique and common entries have been represented in numbers and percentages in brackets. Motif information has been used only while counting.

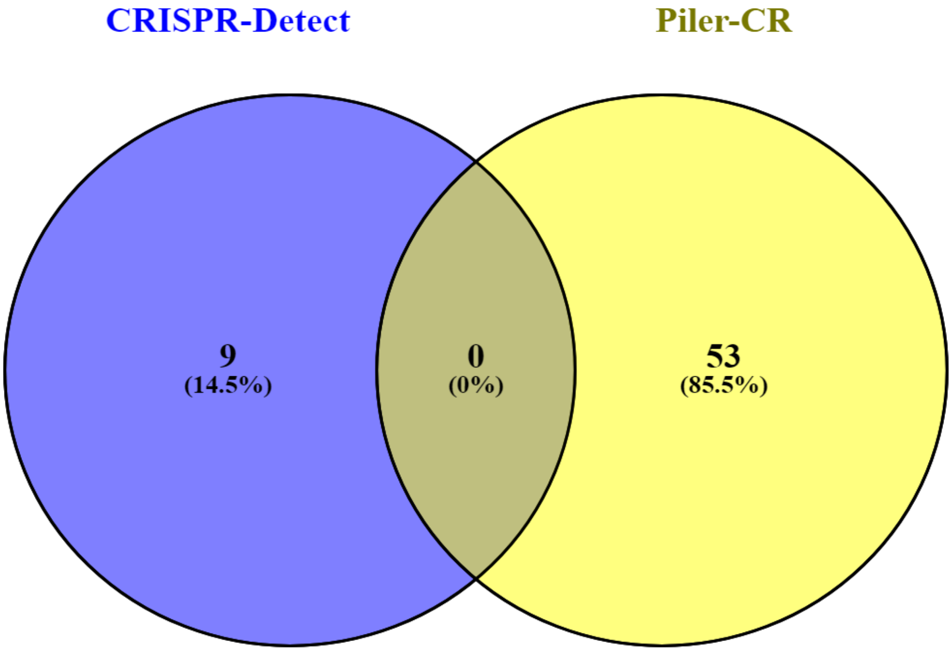


Figure F20: Venn diagram of all CRISPRS extracted by tools CRISPR-Detect (blue) and Piler-CR (yellow). CRT do not report any consensus motif information hence not considered. Unique and common entries have been represented in numbers and percentages in brackets. Motif and repeat length have been used together while counting.

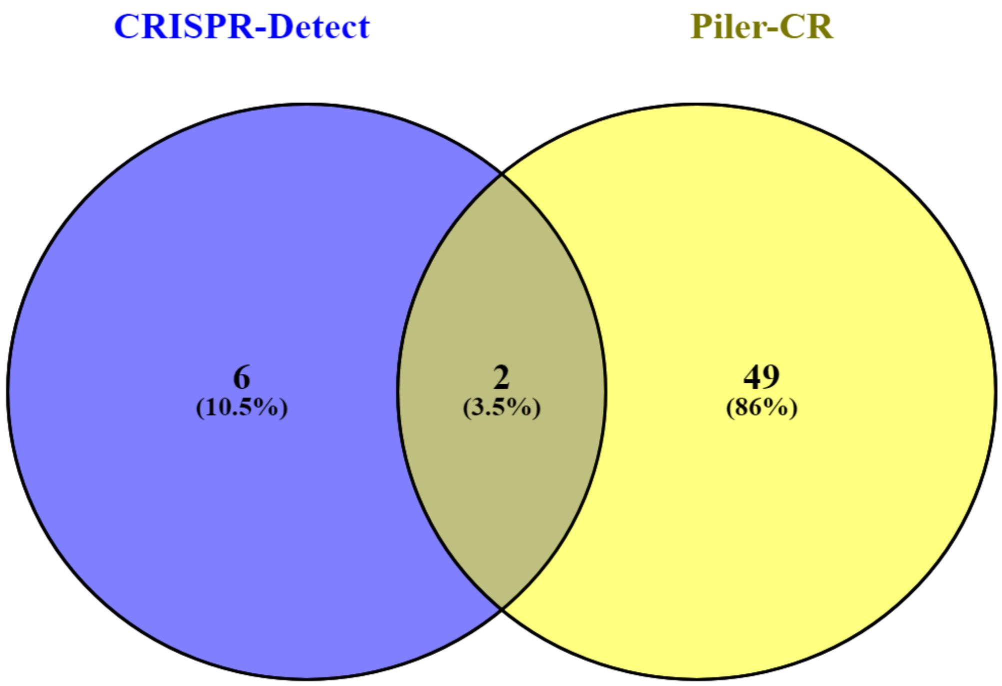


Figure F21: Venn diagram of all CRISPRS extracted by tools CRISPR-Detect (blue) and Piler-CR (yellow). CRT do not report any consensus motif information hence not considered. Unique and common entries have been represented in numbers and percentages in brackets. Motif information has been used only while counting.
